# Genome rearrangements induce biofilm formation in *Escherichia coli* C – an old model organism with a new application in biofilm research

**DOI:** 10.1101/523134

**Authors:** Jarosław E. Król, Donald C. Hall, Sergey Balashov, Steven Pastor, Justin Siebert, Jennifer McCaffrey, Steven Lang, Rachel L. Ehrlich, Joshua Earl, Joshua C. Mell, Ming Xiao, Garth D. Ehrlich

## Abstract

*Escherichia coli* C forms more robust biofilms than the other laboratory strains. Biofilm formation and cell aggregation under a high shear force depends on temperature and salt concentrations. It is the last of five *E. coli* strains (C, K12, B, W, Crooks) designated as safe for laboratory purposes whose genome has not been sequenced. Here we present the complete genomic sequence of this strain in which we utilized both long-read PacBio-based sequencing and high resolution optical mapping to confirm a large inversion in comparison to the other laboratory strains. Notably, DNA sequence comparison revealed the absence of several genes thought to be involved in biofilm formation, including antigen 43, *waaSBOJYZUL* for LPS synthesis, and *cpsB* for curli synthesis. The first main difference we identified that likely affects biofilm formation is the presence of an IS3-like insertion sequence in front of the carbon storage regulator *csrA* gene. This insertion is located 86 bp upstream of the *csrA* start codon inside the −35 region of P4 promoter and blocks the transcription from the sigma^32^ and sigma^70^ promoters P1-P3 located further upstream. The second is the presence of an IS5/IS1182 in front of the *csgD* gene, which may drive its overexpression in biofilm. And finally, *E. coli* C encodes an additional sigma^70^ subunit overexpressed in biofilm and driven by the same IS3-like insertion sequence. Promoter analyses using GFP gene fusions and total expression profiles using RNA-seq analyses comparing planktonic and biofilm envirovars provided insights into understanding this regulatory pathway in *E. coli*.

**IMPORTANCE:** Biofilms are crucial for bacterial survival, adaptation, and dissemination in natural, industrial, and medical environments. Most laboratory strains of *E. coli* grown for decades *in vitro* have evolved and lost their ability to form biofilm, while environmental isolates that can cause infections and diseases are not safe to work with. Here, we show that the historic laboratory strain of *E. coli* C produces a robust biofilm and can be used as a model organism for multicellular bacterial research. Furthermore, we ascertained the full genomic sequence as well as gene expression profiles of both the biofilm and planktonic envirovars of this classic strain, which provide for a base level of characterization and make it useful for many biofilm-based applications.

## Introduction

*Escherichia coli* is a model prokaryote and a key organism for laboratory and industrial applications. *E. coli strain* C was isolated at the Lister Institute and deposited into the National Collection of Type Cultures, London, in 1920 (Strain No. 122). It was characterized as more spherical than other *E. coli* strains and its nuclear matter was shown to be peripherally distributed in the cell (1). *E. coli* C, called a restrictionless strain, is permissive for most coliphages and has been used for such studies since the early 1950’s (2). Genetic tests showed that *E. coli* C forms an O rough R1-type lipopolysaccharide, which serves as a receptor for bacteriophages (3). Its genetic map, which shows similarities to *E. coli* K12, was constructed in 1970 (4). It is the only *E. coli* strain that can utilize the pentitol sugars, ribitol and D-arabitol, and the genes responsible for those processes were acquired by horizontal gene transfer (5). Some research on genes involved in biofilm formation in this strain has been attempted but hasn’t been continued (Federica Briani, Università degli Studi di Milano, personal communication) (6).

Biofilm is the most prevalent form of bacterial life in the natural environment (7-11). However, in laboratory settings, for decades, bacteria have been grown in liquid media in shaking, highly aerated conditions, which select for the planktonic lifestyle. While all laboratory strains of *E. coli*, such as K12, B, W, and Crooks are poor biofilm formers, environmental isolates usually form robust biofilms. These *E. coli* strains can cause diarrhea and kidney failure, while others cause urinary tract infections, chronic sinusitis, respiratory illness and pneumonia, and other illnesses (12-15). Many of these symptoms are correlated with biofilms. A few *E. coli* K12 mutant strains have been described as good biofilm formers, such as the *csrA* mutant or AJW678 (16, 17), but one can claim that these mutants cannot occur in natural conditions. Therefore, it is important to find a safe laboratory strain that can serve as a model for biofilm studies.

We found that the *E. coli* C strain forms a robust biofilm under laboratory conditions. The complete genome sequence of this strain was determined and bioinformatics analyses revealed the molecular foundations underlying this phenotype. Further analyses of differences in gene expression profiles in planktonic and biofilm cells provided an understanding of the regulatory cascades involved in the biofilm process. A combination of experimental and *in silico* analysis methods allowed us to unravel the two major mechanisms that draw the biofilm formation in his strain.

## Results

### Biofilm formation

In our search for a good model biofilm strain, we screened our laboratory collection of *E. coli* strains using the standard 96-well plate assay (18) and the glass slide assay (19) (**Fig. 1A**). We found that the *E. coli* C strain formed robust biofilms on both microscope slides and in 96-well plates. In minimal M9 with glycerol medium, the strain C produced 1.5- to 3-fold more biomass than the other laboratory strains; and in Luria-Bertani (LB) rich medium, the strain C biofilm biomass formation was as much as 7.4-fold higher (**Fig. 1B**).

**Fig. 1.**
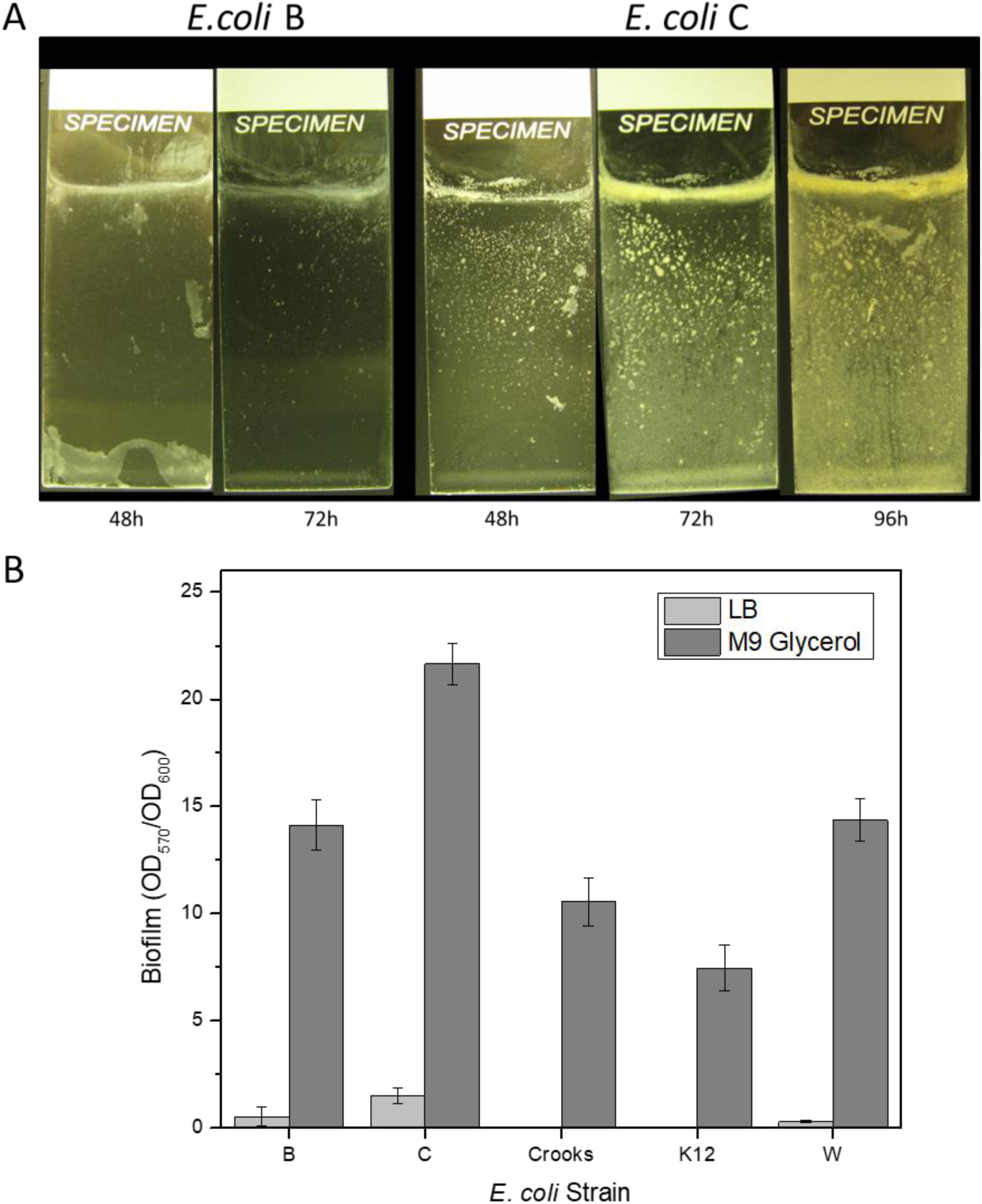
Biofilm formation by *E. coli* strains on (A) microscope slides (LB medium) and (B) 96-well plates (LB and M9 with glycerol).

During the overnight growth in LB medium at 30°C shaken at 250 rpm, we noticed an increased aggregation of bacterial cells in the *E. coli* C culture (**Fig. 2A**). The ratio of planktonic cells to total cells in the culture was 0.35 compared to 0.83 and 0.85 for Crooks and B and 0.98 or almost 1 for K12 and W, respectively (**Fig. 2B**).

**Fig. 2.**
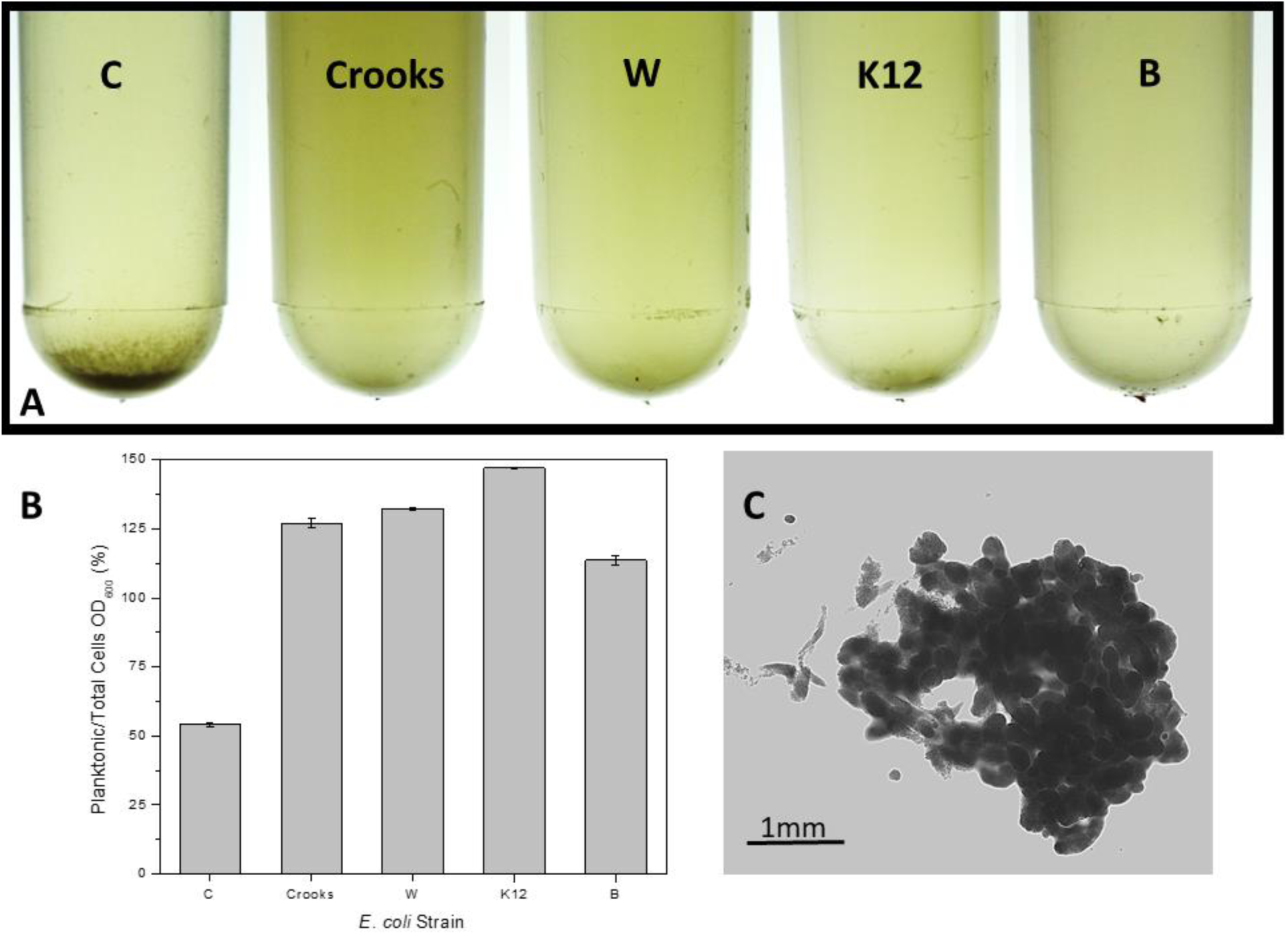
Cell aggregation in overnight culture grown at 30°C in LB Miller broth on shaker at 250 rpm. (**A**) From left: *E. coli* C, *E. coli* Crooks, *E. coli* W, *E. coli* K12 and *E. coli* B. (**B**) Ratio of planktonic cells to total cells measured as OD_600_. (**C**) Microscopic picture of the *E. coli* C precipitate.

Although the aggregation was specific to the *E. coli* C strain, we wanted to test if that precipitation was an active process or just a gravity-driven process. In order to test this hypothesis, the overnight cultures grown in a shaker at 30°C in LB Miller broth were vortexed and aliquoted into standard spectrophotometer cuvettes. Cuvettes were incubated at 12°C, 24°C, and 37°C without shaking and the cell densities (OD_600_) were measured over time. The results showed that despite differences in the initial cell density *E. coli* C strain precipitated less than *E coli* B, K12, and Crooks (**Fig. 3**). The lowest precipitation was observed in the case of strain W (**Fig. 3**). These results showed that the aggregation is an active process related to the bacterial growth under the tested conditions.

**Fig. 3.**
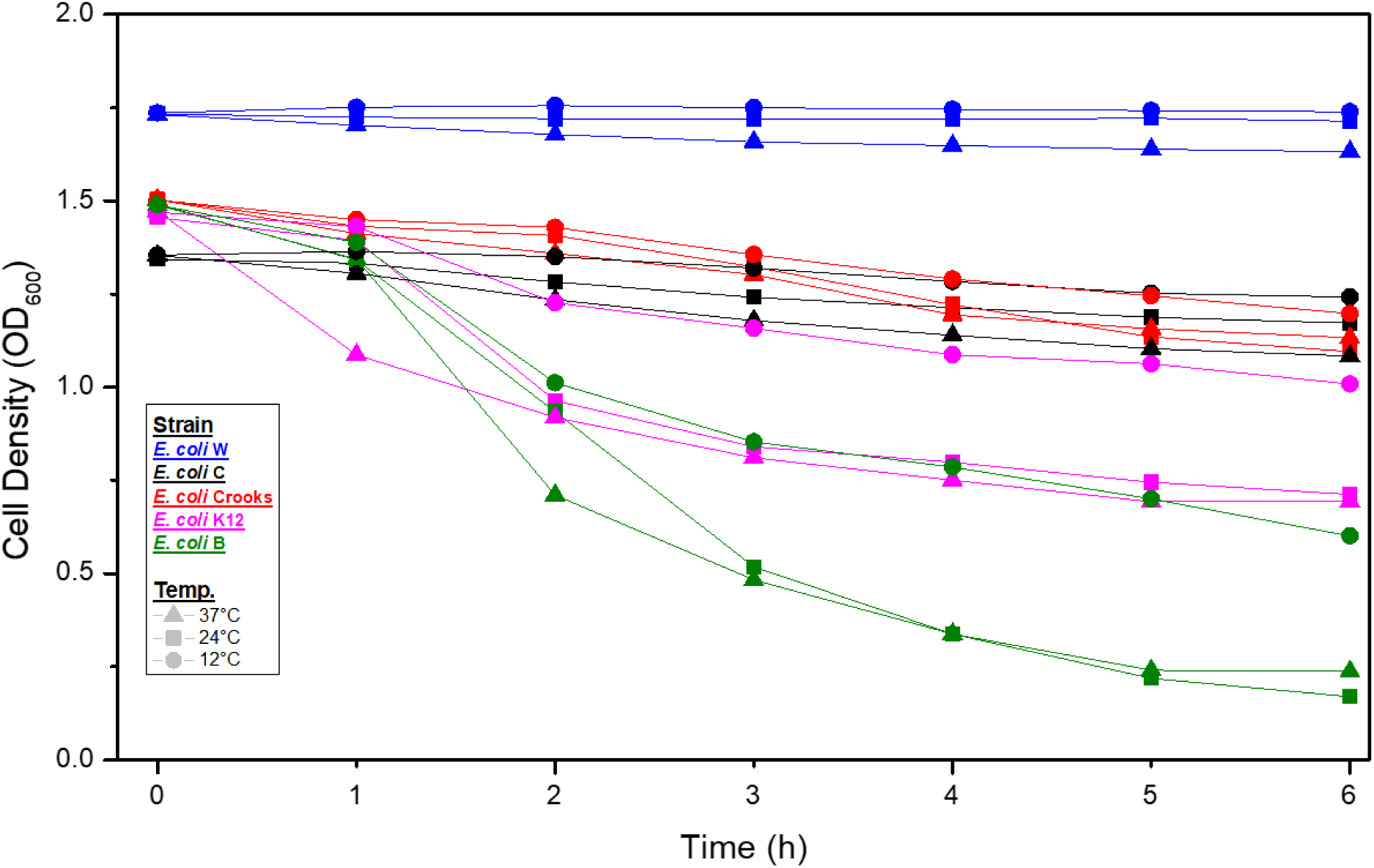
Precipitation of bacterial overnight cultures at 12°C, 24°C, and 37°C without shaking. Bacterial strains are shown in colors, temperatures in shapes (circles 12°C, squares 24°C, and triangles 37°C).

### Precipitation at low temperature depends on salt concentration

Previously, we have described a regulatory loop affecting biofilm formation in a high salt/high pH environment. This loop involved the *nhaR, sdiA, uvrY*, and *hns* genes, as well as the *csrABCD* system (20). We were interested if the precipitation of *E. coli* C depends on NaCl concentration. We grew the bacteria in three LB broth media containing different amounts of salts: Miller broth (1% NaCl), Lennox broth (0.5% NaCl), and a modified Lennox broth with 0.75% NaCl. After overnight growth at 30°C in culture tubes shaken at 250 rpm, we observed a lack of precipitation in standard Lennox medium, while in the modified Lennox medium and Miller broth the ratio of planktonic to total cells was similar (**Fig. 4**). The ratio of planktonic/total cells in the Lennox medium was statistically different (p value < 0.05, Student t-test) from that in media with a higher NaCl concentration and similar to other strains grown in LB Miller broth (**Fig. 2B**).

**Fig. 4.**
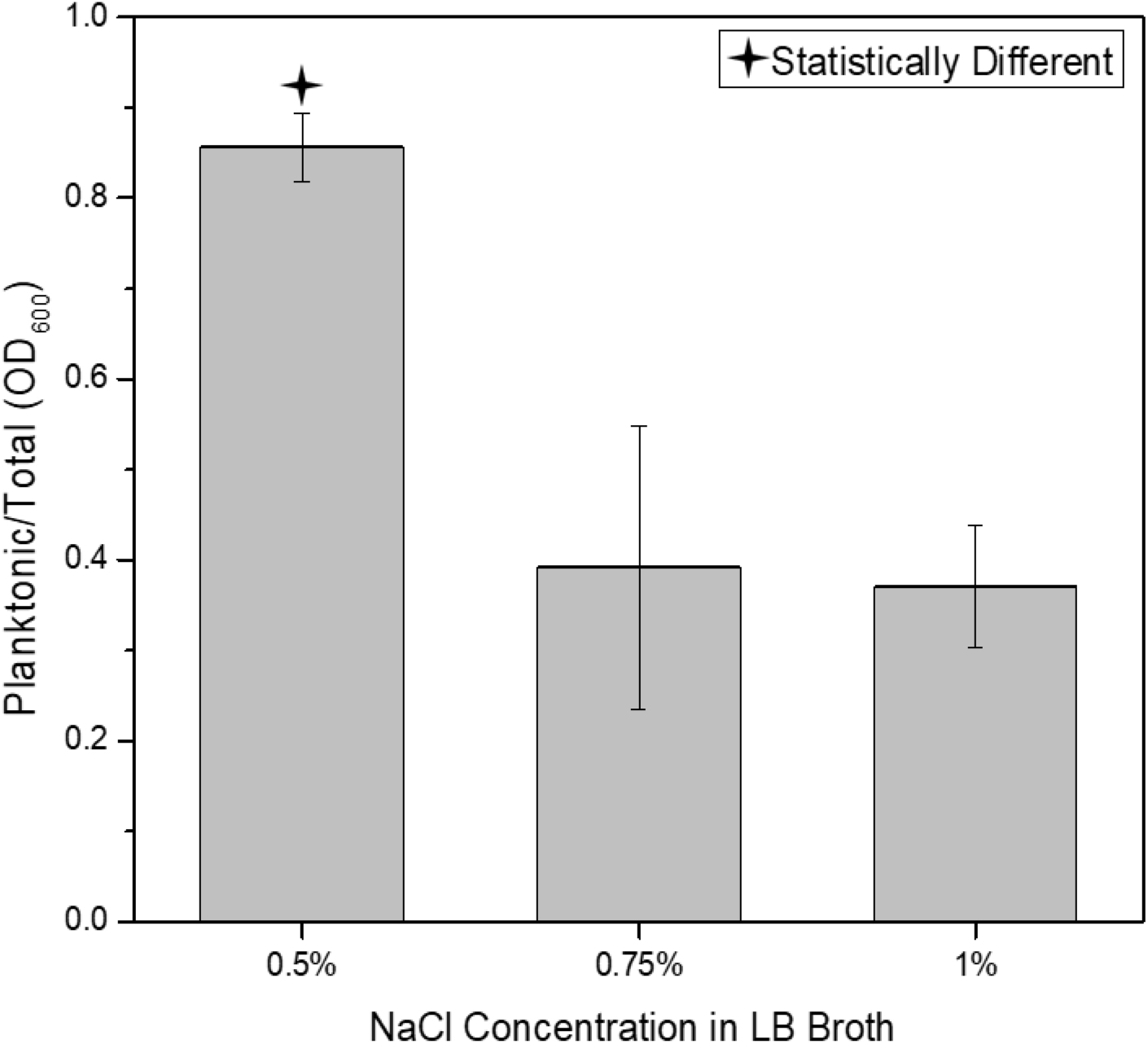
Effect of salt concentration on *E. coli* C precipitation at 30°C.

### *E. coli* C genomic sequence

The genomes of *E. coli* K12, *E. coli* B, *E. coli* W, and *E. coli* Crooks (GenBank:CP000946) have already been sequenced (21-23). To compare the genomic sequences of all five laboratory strains, we sequenced the *E. coli* C genome. The chromosome consisted of 4,617,024 bp and encoded 4,581 CDSs (**Fig. 5**). No extrachromosomal DNA was detected. The mean G+C content was 51%. We identified 7 rRNA operons, 89 tRNA genes, and 12 ncRNAs (total 121 RNA genes CP020543.1).

**Fig. 5.**
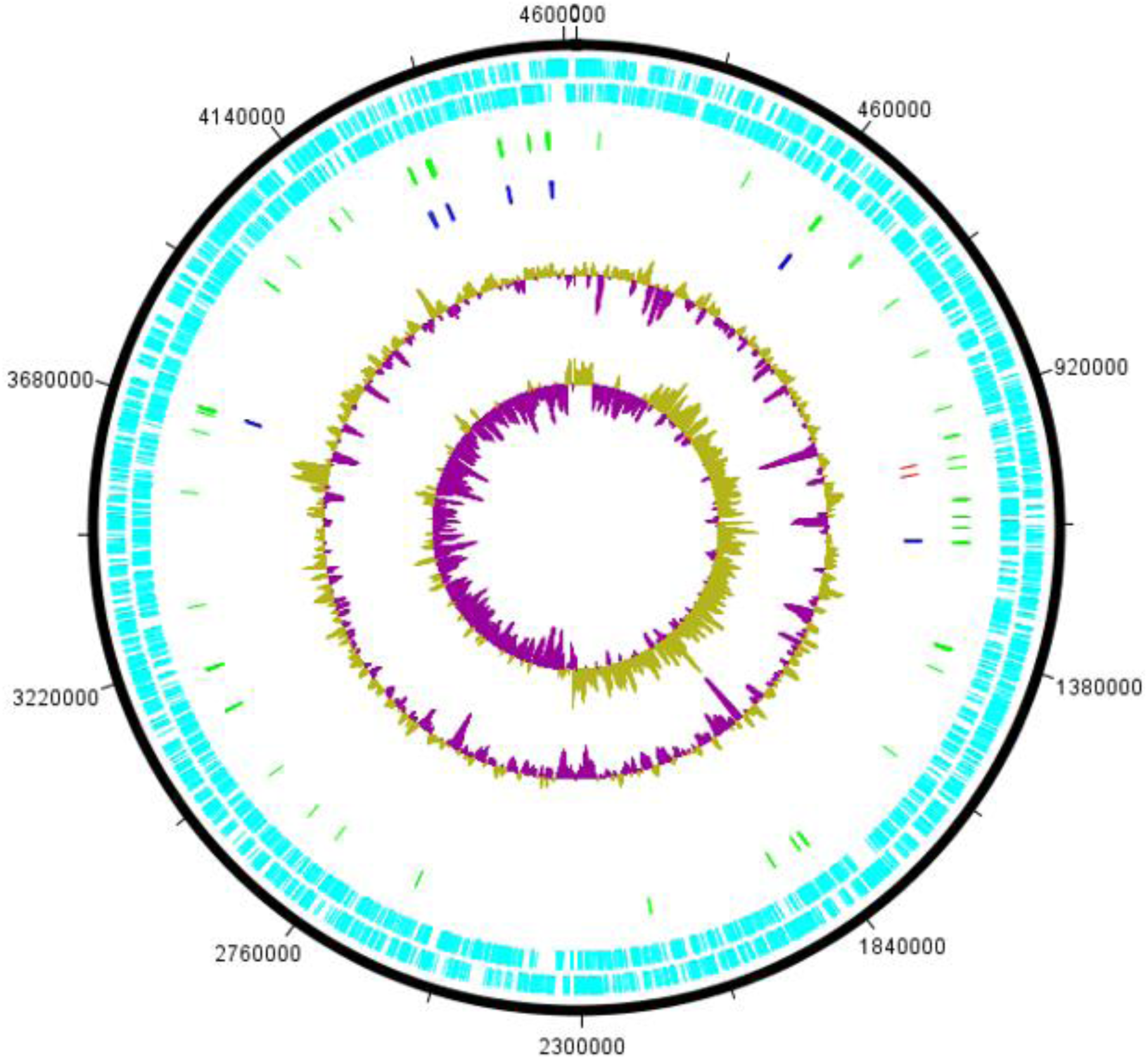
Circular map of the *E. coli* C chromosome (position in bp). The inner circles show GC skew and G+C content. The third circle shows rRNA (blue) and CRISPR (red) clusters. The fourth circle shows hypothetical ORFs (green). Light blue circles represent ORFs on plus and minus strands.

Comparizon with other laboratory *E. coli* strains showed a high degree of synteny except for an inverted 300 kb region between 107 and 407 kb (**Fig. 6**). That inverted region showed also an inverted GC skew in comparizon to the flanking regions, indicating a recent inversion event or an assembly error (**Fig. 5**). To prove that the inversion represented an actual event, we used an optical mapping method (Lam et al. 2012). The order of obtained fluorescently labelled fragments was identical with the *in sillico* constructed map of *E. coli* C chromosome (**Fig. 7A**), indicating the authenticithy of the inversion.

**Fig. 6.**
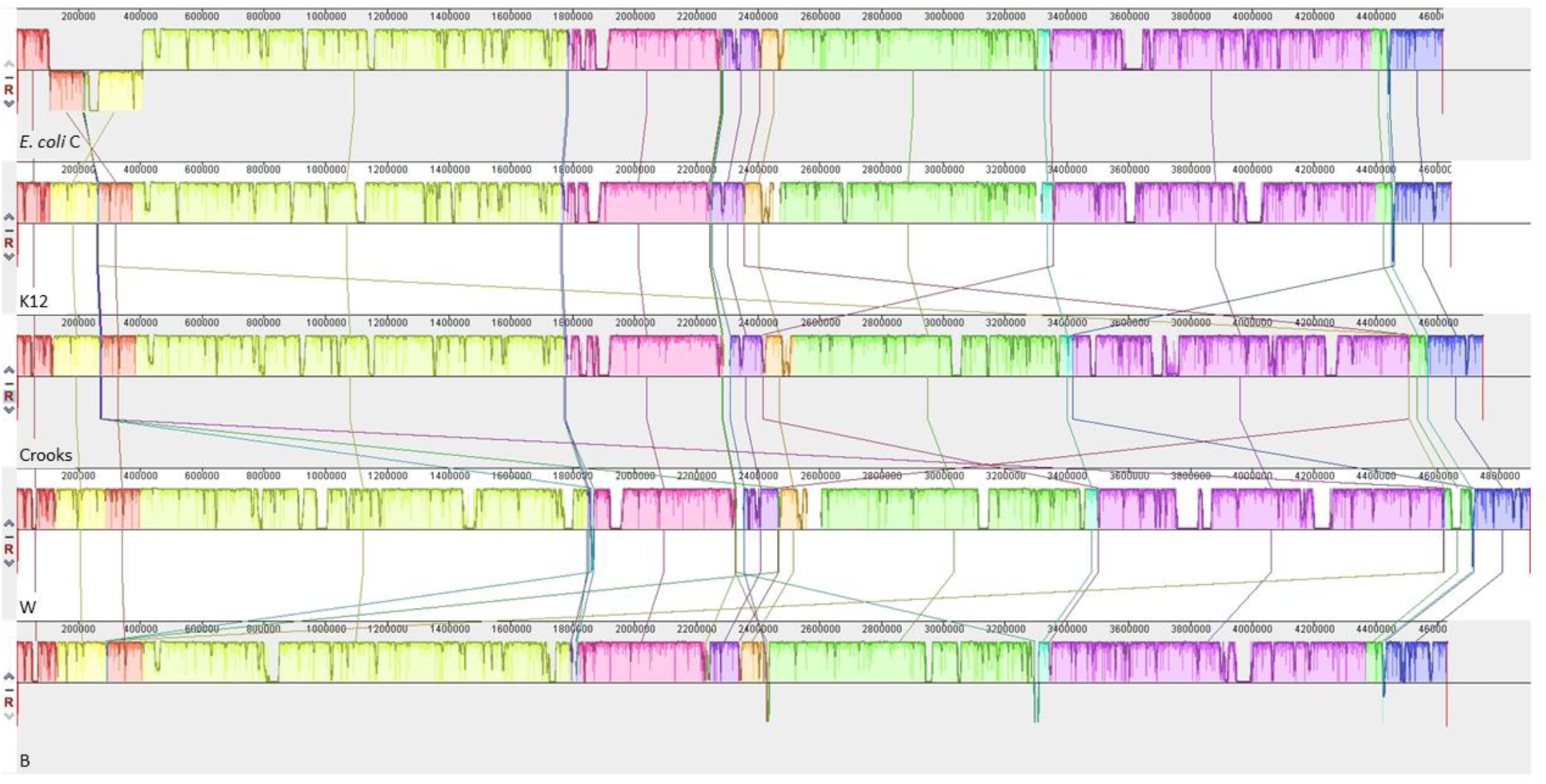
Genome alignment of five *E. coli* strains using Mauve. Each chromosome has been laid out horizontally and homologous blocks in each genome are shown as identically colored regions linked across genomes. The inverted region in *E. coli* C is shifted below the genome’s center axis. From the top: *E. coli* C, K12, Crooks, W, and B.

**Fig. 7.**
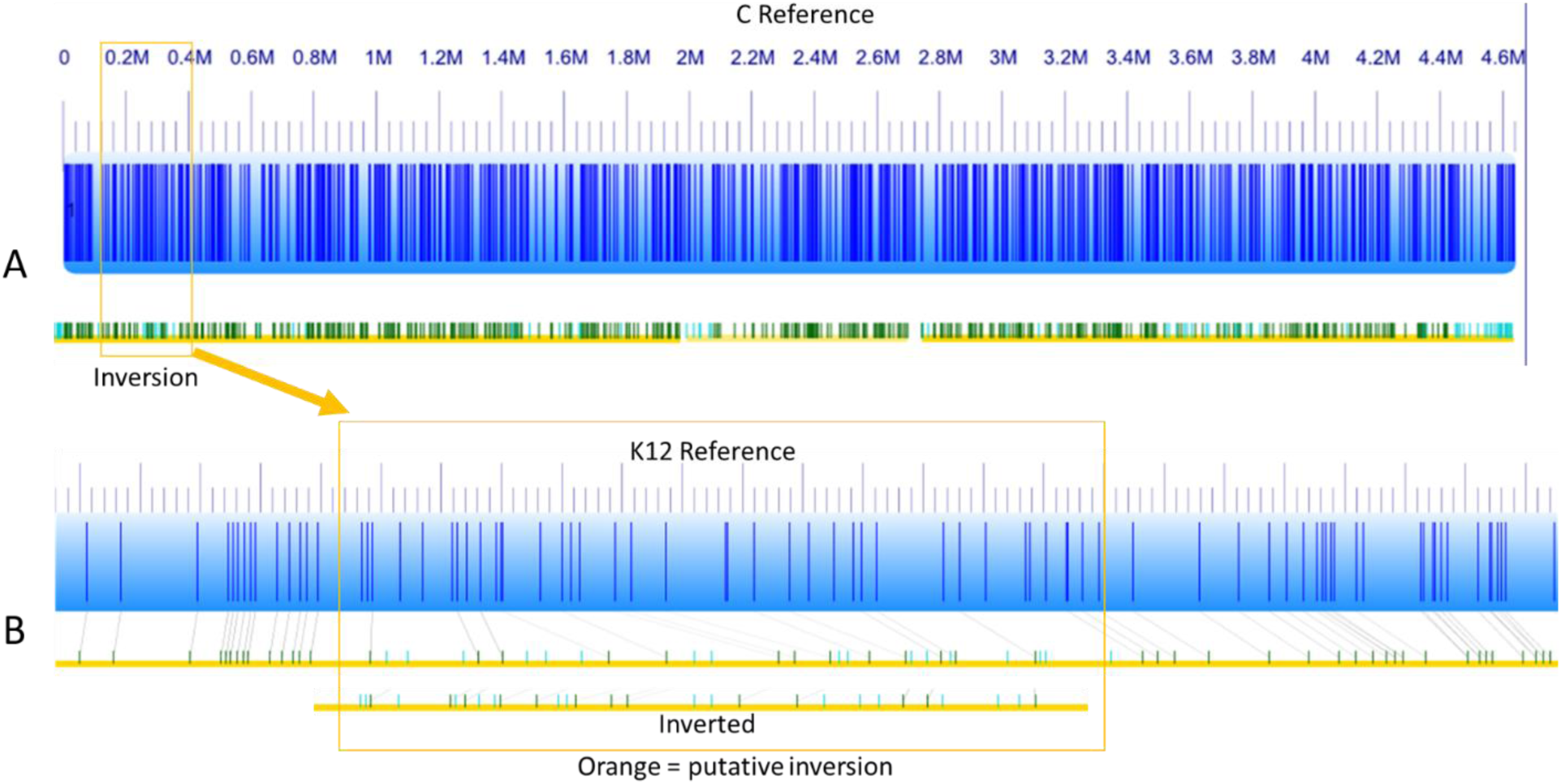
Optical mapping of *E. coli* C chromosome and comparison to K12 strain. (**A**) *In silico* generated map (blue) and optical map (yellow/green) of *E. coli* C. (**B**) *In silico* generated map (blue) and optical map (yellow/green) of *E. coli* K12.

A similar comparizon of *E. coli* K12 maps confirmed the stringency and precision of the optical mapping results (**Fig. 6B**). Comparizon between the two optical maps confirmed that the PacBio-predicted inversion of the 300-kb DNA fragment was indeed a real event (**Fig. 6B**).

### Genetic content

A neighbor joining tree based on gene possession showed that *E. coli* C was most similar to the K12 strain (**S1 Fig**). A comparison of chromosomal protein-coding orthologs among the laboratory strains showed that, out of the 5,686 predicted CDSs, 3,603 were shared among all five strains. Only 33 genes were present in all four of the other lab strains that were absent in E. *coli* C (S1 Tab) (**Fig. 8**). Out of 177 genes that were unique to *E. coli C*, 108 encoded transposases or unknown proteins and 69 CDSs showed homology to known proteins (**Tab. 1**).

**Fig. 8.**
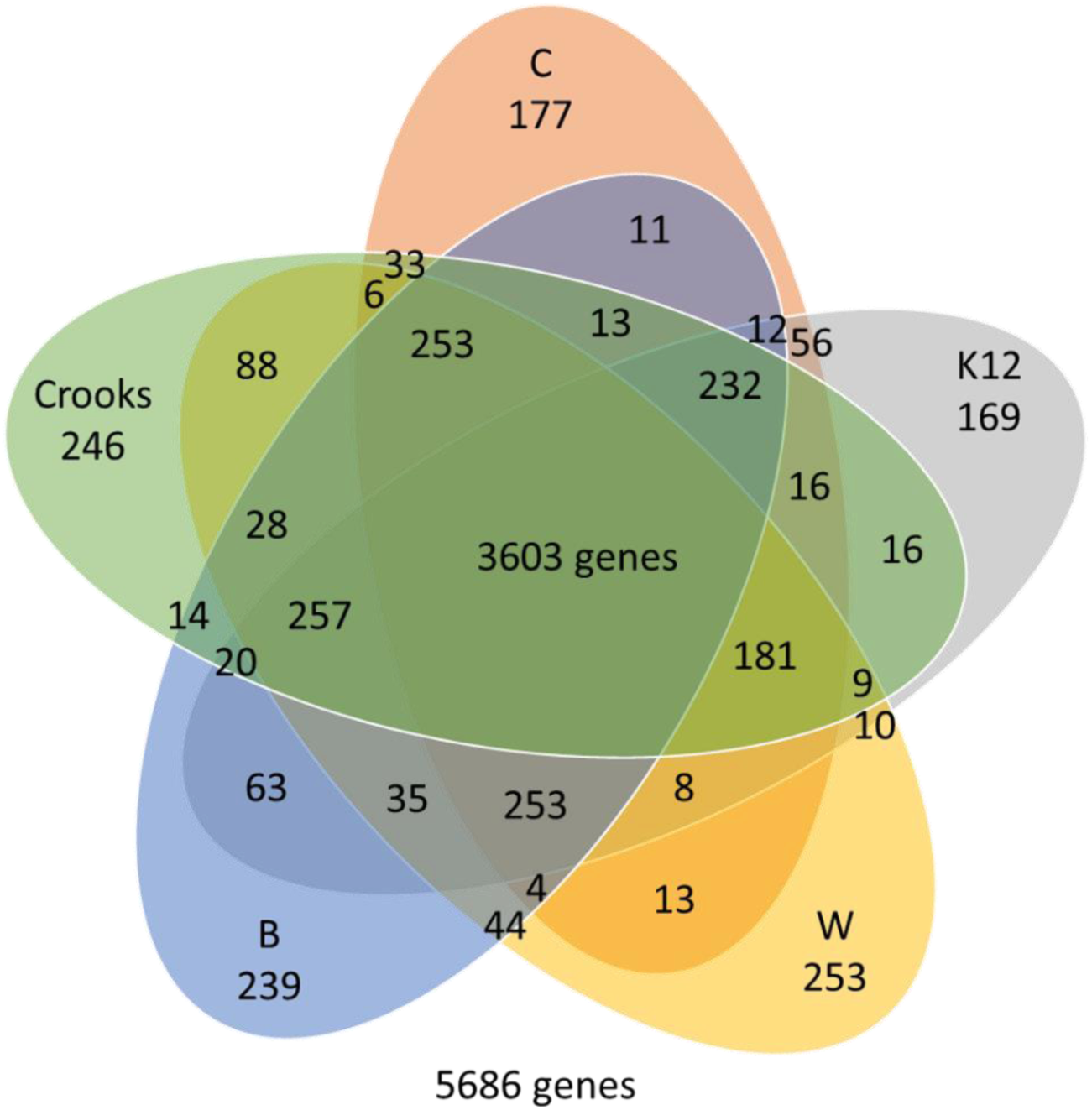
Comparison of orthologous CDSs among C, W, K-12, B, and Crooks strains. The number of shared genes, the number of unique genes, and the genes shared between one, two, three, and four strains are shown.

### Genes involved in biofilm formation

Several genes have been ascribed active roles in biofilm formation in *E. coli* (24-26). One of the most important is the *flu* gene encoding the antigen 43 protein (27). In liquid culture, Ag43 leads to autoaggregation and clump formation rapidly followed by bacterial sedimentation. Surprisingly, the *flu* gene was not present in the *E. coli* C genome. We identified a few autotransporter encoding genes, which showed partial homology to Ag43, such as B6N50_05815 (50% similarity over 381aa); however, the homology was too weak to suggest that these genes could play a similar role.

Surface polysaccharides often play an important role in biofilm formation (25, 28). *E. coli* C forms an O rough R1-type lipopolysaccharide, which serves as a receptor for bacteriophages (Feige and Stirm 1976). Out of the 14 *waa* genes present in *E. coli* K12, we were able to find only 6 in *E. coli* C. Out of 5 genes *waaA, waaC, waaQ, waaP*, and *waaY*, which are highly conserved and responsible for assembly and phosphorylation of the inner-core region (29), only the first 4 were present in *E. coli* C (**S2 Fig**). Two remaining genes in *E. coli* C were *waaG*, whose product is an α-glucosyltransferase that adds the first residue (HexI) of the outer core, and *waaF*, which encodes for a HepII transferase (29). Biofilm formation by a deep rough LPS *hldE* mutant of *E. coli* BW25113 strain was strongly enhanced in comparison with the parental strain and other LPS mutants. The *hldE* strain also showed a phenotype of increased autoaggregation and stronger cell surface hydrophobicity compared to the wild-type (30). The gene *hldE*, which encodes for a HepI transferase, was found in the *E. coli* C strain. Other mutants in LPS core biosynthesis, which resulted in a deep rough LPS, have been described to decrease adhesion to the abiotic surfaces (31); therefore, we assumed that other genes in this family would not be responsible for the increased biofilm formation by *E. coli* C.

We noticed also that *wzzB*, a regulator of length of O-antigen component of lipopolysaccharide chains was mutated by an IS3 insertion. Another IS insertion was located in UDP-glucose 6-dehydrogenase (B6N50_08940). Both these genes were located at the end of a long 35 operon-like gene stretch in *E. coli* C, including *wca* operon (32) consisting of 19 genes involved in colanic acid synthesis.

We found that the region involved in biosynthesis of poly-β-1,6-N-acetyl-glucosamine (PGA) was almost 100% identical in both K12 and C strains.

Other types of structures involved in biofilm formation are fimbriae, curli, and conjugative pili (24, 25, 33). Type 1 pili can adhere to a variety of receptors on eukaryotic cell surfaces. They are well-documented virulence factors in pathogenic *E. coli* and are critical for biofilm formation on abiotic surfaces (34-38). Type 1 pili are encoded by a contiguous DNA segment, labeled the *fim* operon, which contains 9 genes necessary for their synthesis, assembly, and regulation (39, 40). In *E. coli* C, almost the entire *fim* operon except the *fimH*, which codes for the mannose-specific adhesin located at the tip of the pilus, was absent and replaced by a type II group integron (**S3 Fig**). The entire *fin* operon is driven by a single promoter located upstream of the *fimA* gene; therefore, it is possible that the *fimH* gene is not expressed in *E. coli* C. Although we cannot exclude the role of FimH in autoaggregation of *E. coli* C, reports that the function of FimH was inhibited by growth at temperatures at or below 30°C (41) make it highly unlikely.

Chaperone-usher (CU) fimbriae are adhesive surface organelles common to many gram-negative bacteria. *E. coli* genomes contain a large variety of characterized and putative CU fimbrial operons (42). Korea at al. characterized the *ycb, ybg, yfc, yad, yra, sfm*, and *yeh* operons of *E. coli* K-12, which display sequence and organizational homologies to type 1 fimbriae exported by the CU pathway (43). They showed that, although these operons were poorly expressed under laboratory conditions, 6 of them were nevertheless functional when expressed and promote adhesion to abiotic and/or epithelial cell surfaces (43). A total of 10 CU operons have been identified in *E. coli* K12 MG1655 (42). We identified all 10 CU operons in the *E. coli* C genome. Furthermore, we found that the IS5 insertion in the K12 *yhcE* gene was not present in *E. coli* C (**S4A Fig**). We also noticed that two insertion sequences were inserted in the *yad* region (**S4B Fig**).

Curli are another proteinaceous extracellular fiber involved in surface and cell-cell contacts that promote community behavior and host cell colonization (44). Curli synthesis and transport are controlled by two operons, *csgBAC* and *csgDEFG*. The *csgBA* operon encodes the major structural subunit CsgA and the nucleator protein CsgB (45). CsgC plays a role in the extracellular assembly of CsgA. In the absence of CsgB, curli are not assembled and the major subunit protein, CsgA, is secreted from the cell in an unpolymerized form (44). The *csgDEFG* operon encodes 4 accessory proteins required for curli assembly. CsgD is a positive transcriptional regulator of the *csgBA* operon (45). We found that the intergenic region between *csgBA* and *csgDEFG* has been modified in *E. coli* C. An IS5/IS1182 family transposase was inserted between 106 bp upstream of the *csgD* gene and 96 bp inside the *csgA* gene (**S5 Fig**). The entire *csgB* gene as well as the first 32aa of CsgA have been deleted. The full CsgA protein in *E. coli* K12 contains 151aa while the truncated version in strain C consisted of only 107aa and might not be expressed. Furthermore, *csgD* expression is driven by a promoter located ∼130 bp upstream (46, 47). The IS5/IS1182 family transposase inserted between that promoter and the *csgD* gene was transcribed in the same direction, so it might not cause a polar mutation but definitely would interfere with the sophisticated regulation of *csgD* expression by multiple transcription factors (46, 47). As *E. coli* C did not carry any extrachromosomal DNA, conjugative pili, which usually play an important role in biofilm formation (48), were not analyzed.

Biofilm formation is a bacterial response to stressful environmental conditions (9). This response requires an orchestra of sensors and regulators during each step of the biofilm formation process. We analyzed a few of the most important mechanisms, such as CpxAR, RcsCD, and EnvZ/OmpR (25). In all three cases, we observed the same gene structure and a high degree of DNA sequence identity between *E. coli* C and K12 strains.

Another regulatory loop includes the carbon storage regulator *csrA* and its small RNAs (49). Mutations within the *csrA* gene induced biofilm formation in many bacteria (17, 49). Recently, the CsrA regulation has been connected with multiple other transcription factors, including NhaR, UvrY, SdiA, RecA, LexA, Hns, and many more (20, 50). The regulatory loop with NhaR protein drew our attention as it is responsible for integrating the stress associated with high salt/high pH and low temperature (20). We found that the *nhaAR* and *sdiA*/*uvrY* regions of *E. coli* C were almost identical with the corresponding regions in the K12 strain. We amplified and sequenced the *csrA* gene from the *E. coli* C strain to verify its presence and integrity (**S6 Fig**). Detailed analysis of the *csrA* region revealed the presence of an IS3-like insertion sequence 86 bp upstream of the ATG codon (**Fig. 9**). The *csrA* gene is driven by 5 different promoters (51). The distal (−227 bp) promoter P1 is recognized by sigma^70^ and sigma^32^ factors and enhanced by DskA. The P2 (−224 bp) promoter depends on sigma^70^. Both P1 and P2 promoters are relatively weak promoters (51). The P3 promoter is located 127 bp upstream of *csrA* and it is recognized by the stationary RpoS (sigma^32^) polymerase. This promoter is the strongest promoter of *csrA* gene. Promoters P4 and P5 are located 52 bp and 43 bp, respectively, upstream of the *csrA* gene. These promoters are driven by the sigma^70^ polymerase and are active mainly during exponential growth (51). The IS3 insertion was located within the −35 region of the P4 promoter. That location should almost completely abolish expression of the *csrA* gene in the stationary phase of bacterial growth and probably was the main reason for increased biofilm production by the *E. coli* C strain. Both small RNAs, *csrB* and *csrC*, which regulate CsrA activity, were found unchanged in the *E. coli* C genome.

**Fig. 9.**
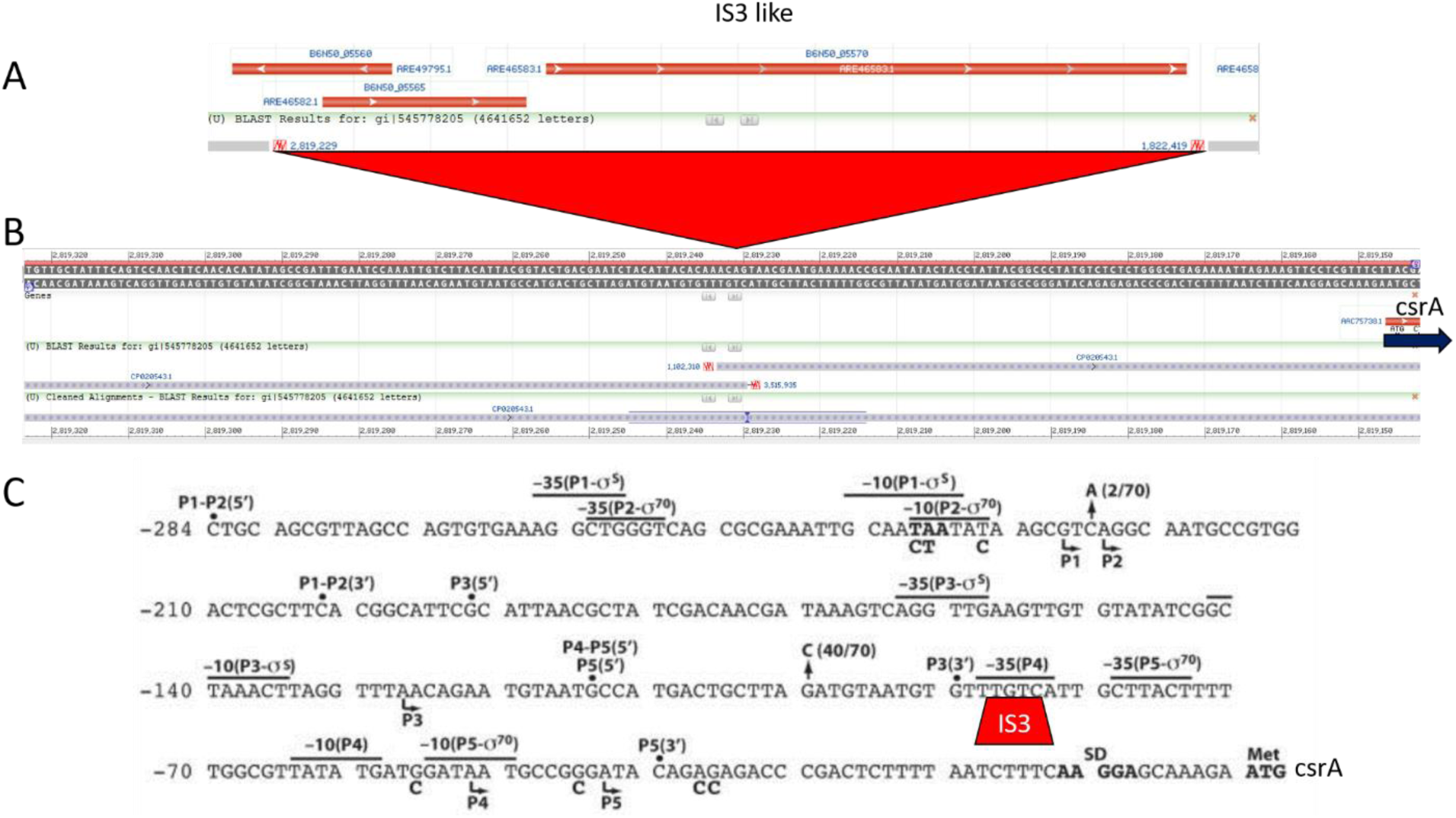
Insertion of IS3 like sequence in the promoter region of *csrA* gene. (**A**) Structure of the IS3 like sequence; (**B**) genome view of K12 (upper) *csrA* promoter region blast results with *E. coli* C; (**C**) *csrA* promoter region (51) with the IS3 insertion site.

Sequence homology search of GenBank available *E. coli* genomes revealed that identical *csrA* promoter regions were present also in strains WG5 (CP024090.1) and NTCT122 (LT906474.1), while some other strains contained the IS3 sequence but missed the upstream region (data not shown).

### Confirmation of IS3 insertion and its complementation by overexpression of *csrA* gene

First we compared the biofilm formation ability of *E. coli* C and the K12 *csrA* mutant. The 72-hour-old biofilms of both strains formed on microscope slides were similar (**S7A Fig**). The 24-hour 96-well plate biofilm assay showed that at 37°C the K12 *csrA* mutant formed 30% more biofilm than *E. coli* C (p value = 0.001, Student t-test). At 30°C strain C produced more biofilm, but the difference was not statistically significant (**S7B Fig**). To confirm the presence of the IS3 insertion in the *csrA* promoter region, we designed PCR primers specific for the *alaS*-*csrA* intergenic region. Amplification results confirmed the presence of IS3 in the *E. coli* C promoter region (**S8 Fig**).

To see if extrachromosomal expression of the CsrA protein affects the precipitation phenotype, we cloned the *csrA* gene downstream of a *plac* promoter in pBBR1MCS-5 (52), resulting in plasmid pJEK718 or downstream of the constitutive *pcat* (chloramphenicol) promoter in pJEK786. Plasmids were transformed into *E. coli* C strain and the resulting clones were grown in LB Miller broth (30°C, 250 rpm). The results showed that the ratio of planktonic to total cells in *E. coli* C carrying both constructs overexpressing the *csrA* gene was ∼1.8 times higher (p value > 0.0001, Student t-test) than in the control carrying the non-recombined vector (**Fig. 10**). We also noticed that the control strain showed a slightly higher amount of planktonic cells than the plasmidless control (0.46 vs. 0.36), although the difference was not statistically significant (p value = 0.09, Student t-test).

**Fig. 10.**
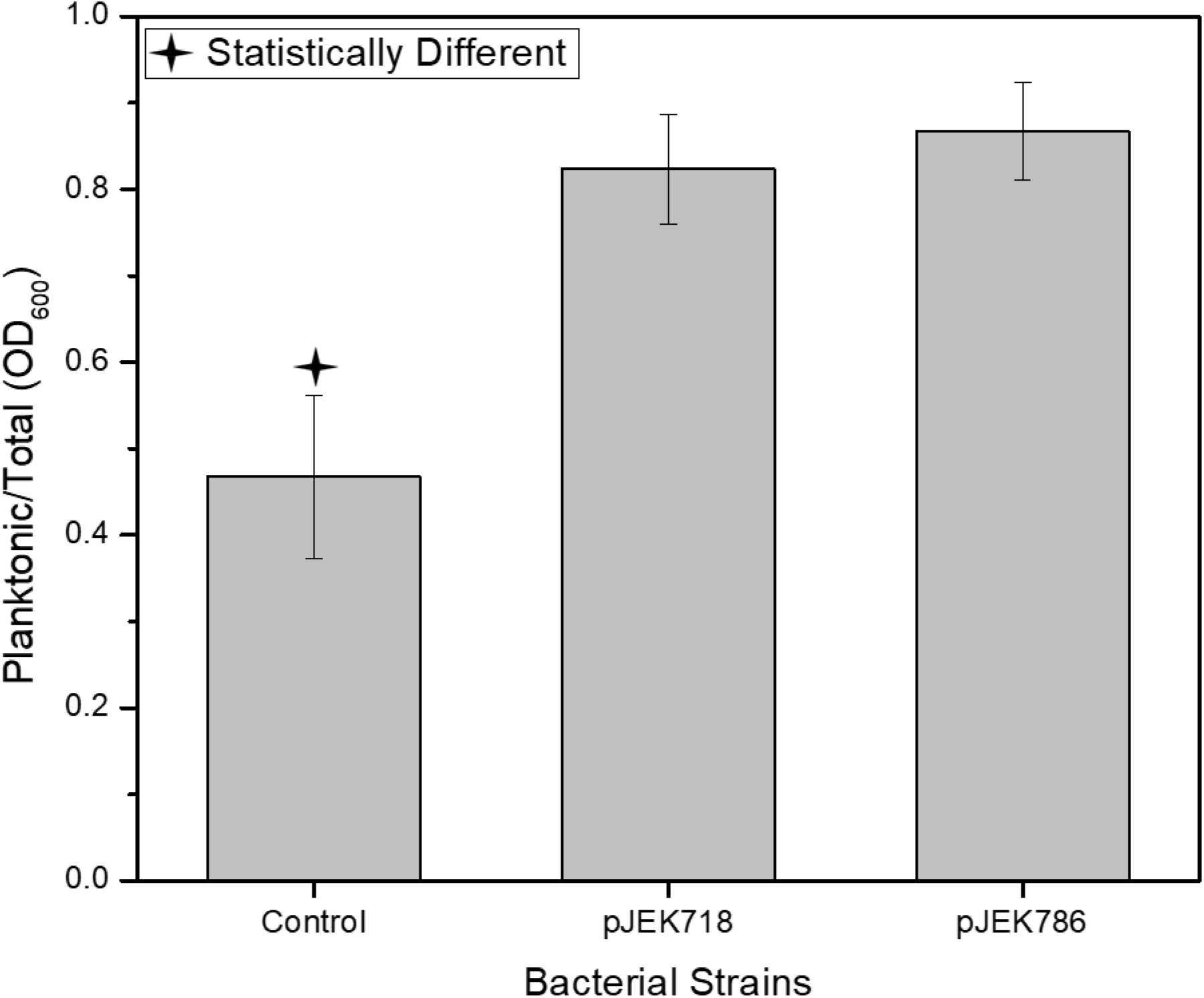
Complementation of *E. coli* C aggregation phenotype by introduction of pJEK718 and pJEK786 plasmids overexpressing the CsrA protein.

### Differential gene expression in biofilm and planktonic cells

In order to identify the genes involved in biofilm formation, we investigated the global transcriptional differences between mature *E. coli* C biofilms and planktonic cultures. Previously, differences in the expression profiles between *E. coli* biofilm and planktonic cells were done using microarrays, which were state of art at the time (53-55) (56). Since that time, only one report has been published on *E. coli* biofilm expression profiles in aerobic and anaerobic biofilms and planktonic cells using a high-throughput RNA-seq technology (57). We used RNA-seq technology to analyze *E. coli* C expression profiles. Total RNA was extracted from a 5-day-old slide biofilms which were transferred to a fresh LB Miller broth daily. As planktonic cells we used cells detached from the biofilm from the last (5^th^ day) passage. Three biological replicates were analyzed.

In general, we found that the expression of 4,124 genes from the total number of 4,702 genes was either not changed or not detected under either condition. Four hundred two genes were overexpressed in biofilm and 177 genes showed an increased expression in planktonic cells (**Fig. 11A, B**).

**Fig. 11.**
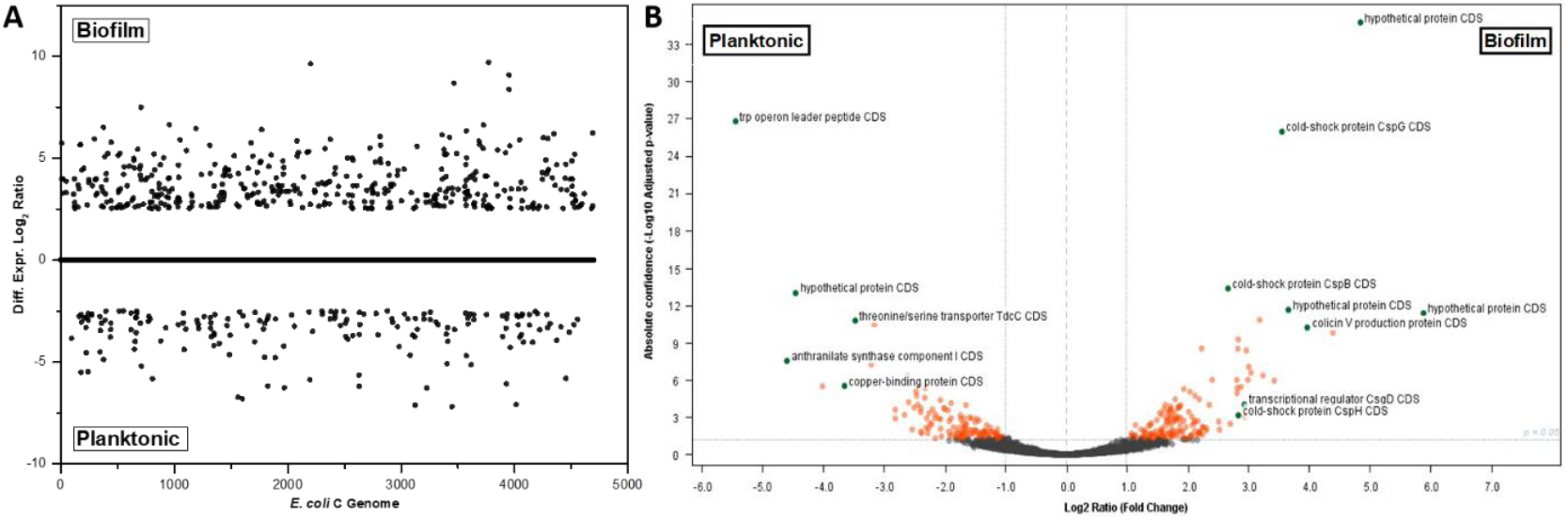
Overview of the differential gene expression among the *E. coli* C envirovars. (**A**) Distribution of genes along the chromosome; only genes significantly altered in expression (average fold change ≥ [2.0]) were indicated. (**B**) Volcano plot with differential expression and statistical significance.

Among the top 80 differentially expressed genes between biofilm and planktonic cells (**Tab. 2**), we did not observe any obvious candidates for being biofilm determinant genes in *E. coli* C. However, it was interesting that, even when biofilms were grown at 37°C, several of the most highly differentially expressed genes were 5 different cold shock proteins, which did not belong to the same transcription units. This was not surprising, as previously the CspD gene has also been linked to biofilms and persister cell formation (58) and other Csp proteins were also involved in biofilm formation (59).

Another highly expressed biofilm gene encoded for an *acrEF/envCD* operon transcriptional regulator (B6N50_02560). One report suggested that the *acrEF/envCD* operon transcriptional regulator interacts with one of the CU type putative fimbrial adhesin YraK (60). We noticed that the expression of the *yraK* gene was slightly increased in biofilm (Diff.Expr.Log2 Ratio 1.6, p value = 0.15) while other genes encoding YraCU were downregulated (**S2 Tab**).

An additional interesting observation made based on the expression profile was that, even though the *fim* operon with its promoter region was deleted in *E. coli* C, the remaining *fimH* gene was hugely overexpressed in biofilm (Diff.Expr.Log2 Ratio = 5.18, p value = 0.08) (**S2 Tab**), indicating the presence of a previously uncharacterized promoter in front of this gene. A similar situation was observed in the case of the *csgD* regulator. Although the entire *csgD* promoter has been deleted in the *E. coli* C, its expression was still much higher in biofilm than in planktonic cells (Diff.Expr.Log2 Ratio = 3.48, p value = 3.87×10^−05^) (**S2 Tab, Fig. 11B**), suggesting that the remaining ∼100-bp upstream region of that gene was enough to drive its expression at a highly elevated level or the existence of an additional promoter.

As we had speculated that the biofilm phenotype of the *E. coli* C strain depends on the *csrA* gene, we were interested in its expression. We found that the *csrA* was upregulated in the biofilm compared to the planktonic cells (Diff.Expr.Log2 Ratio = 1.79, p value = 0.01) (**S2 Tab**). The observed *E.coli* C *csrA* expression was rather unexpected, as higher CsrA levels have been associated with increased motility and decreased biofilm formation (49). The global effect of the CsrA protein on *E. coli* gene expression has been studied in great detail by the Romeo and Babitzke groups (49, 50, 61). However, all of their experiments were conducted using planktonic cells, and so a direct comparison with our data would be irrelevant. We reached a similar conclusion when we tried to compare expression profiles from microarray tests. Unfortunately, the most recent biofilm RNA-seq experiment was also conducted using M9 minimal medium, also making data comparison irrelevant (57).

### Expression of *csrA* promoter in *E. coli* K12 and *E. coli* C

To analyze activities of the *csrA* promoter from *E. coli* C, we cloned PCR products containing sequences upstream of the *csrA* gene (**S8 Fig**) into a pAG136 plasmid vector carrying promoterless EGFP-YFAST reporters (pJEKd1750) (62). The *E. coli* C *csrA* promoter was overexpressed in both strains, however the promoter activity was much stronger in the native strain then in K12 (**S9 Fig**). We notice that the highest differences (3.2 and 2.4, at 37°C and 30°C, respectively) occurred at the late exponential phase (∼4.5h and ∼10h) (**S9 Fig**). We noticed that the presence of an additional copy of *pcsrA* in a high copy number plasmid induced precipitation of *E. coli* C at 37°C. The ratio of planktonic/total cells was similar (**S10 Fig**) to that obtained for the parental *E. coli* C strain at 30°C (**Fig. 2** and **Fig. 4**) (0.36 and 0.35, respectively).

The aggregation phenotype was correlated with the highest *pcsrA* activity at the entrance to the stationary phase (data not shown). As the aggregation might affect the measurements we decided to use a colony assay to measure the promoter activity over the long time. The LB agar plates with spots of *E. coli* C and K12 carrying pJEKd1751 reporter plasmids with a short half-life form of GFP[ASV] were incubated at 30°C and 37°C and the fluorescence activity was measured by a Typhoon 9400 Variable Mode Imager (**Fig. 12**). The data showed an increased *pcsrA* activity over the 72h time period in both strains with much higher activity in the native *E. coli* C strain (**Fig. 12**). The highest differences between both strains, 8.15 and 4.71 were observed at 72h at 30°C and 37°C, respectively (**Fig. 12**). As the half-life of the GFP[ASV] is only 110 min.(63), we concluded that in the K12 strain *pcsrA* promoter was active mostly at the stationary phase while in the *E. coli* C its activity was quasi constitutive, but also enhanced at the stationary phase (**Fig. 12**). To test that hypothesis we analyze the spatial expression of the pcsrA promoter in 72h old bacterial colonies using a fluorescence microscope (**Fig. 13**). The pictures fully supported our premises. In the *E. coli* C the entire colony showed an intensive fluorescence with the highest level in the center (**Fig. 13A**). In the K12 strain we noticed 5 discrete zones with different fluorescence activities (**Fig. B**). The edge of the colony which should consist of the youngest, still dividing and metabolically active cells showed the lowest, while the center of the colony with the oldest cells showed the highest fluorescence (**Fig. 13B**).

**Fig. 12.**
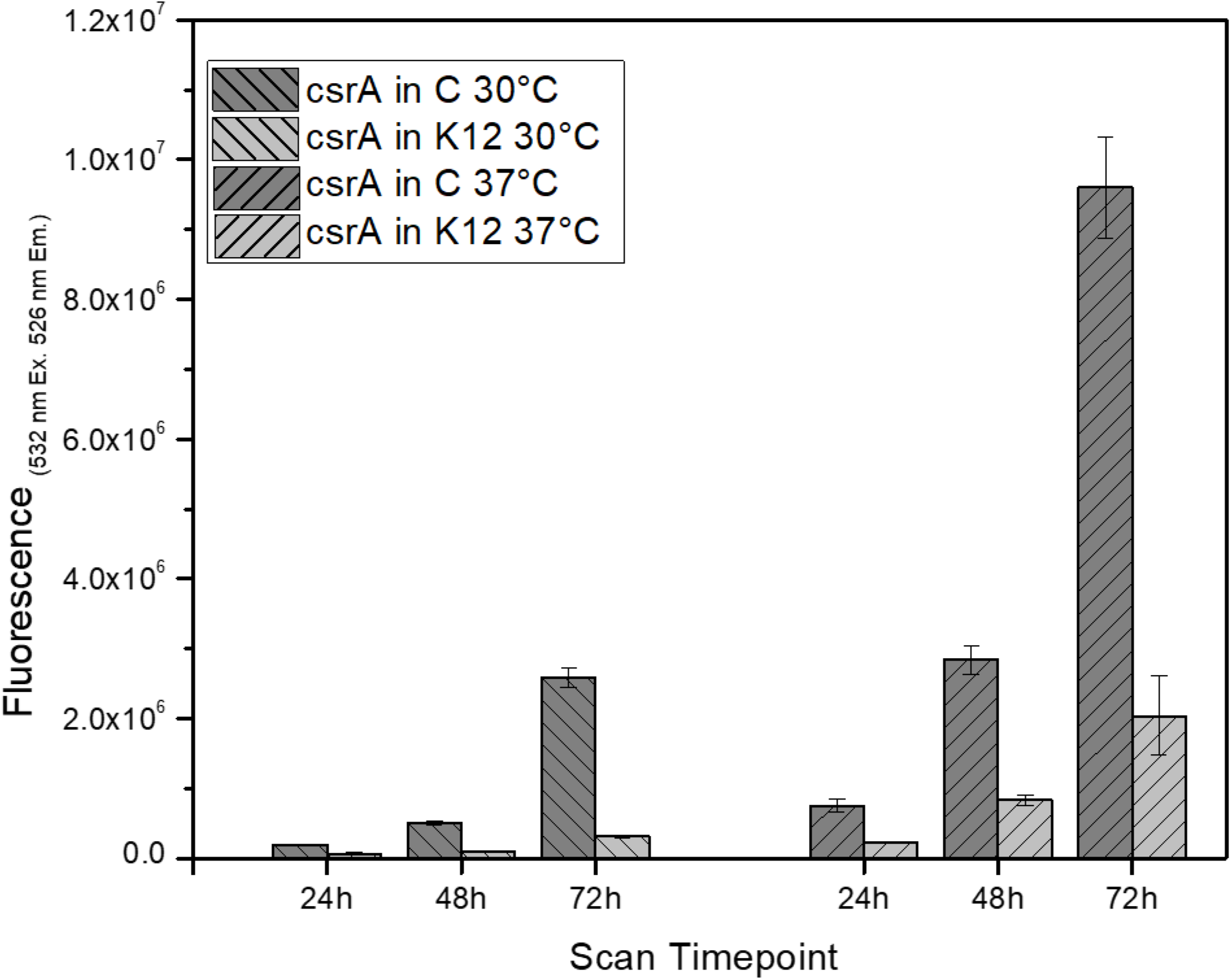
Activity of *pcsrA* promoter (pJEKd1571) in 24h, 48h and 72h old colonies of *E. coli* C and K12 grown at 30°C and 37°C on LB Miller agar plates. Data represents the mean values from 3 biological replicates each containing 3 colonies. Differences between strains at all time points and conditions were statistically significant.

**Fig. 13.**
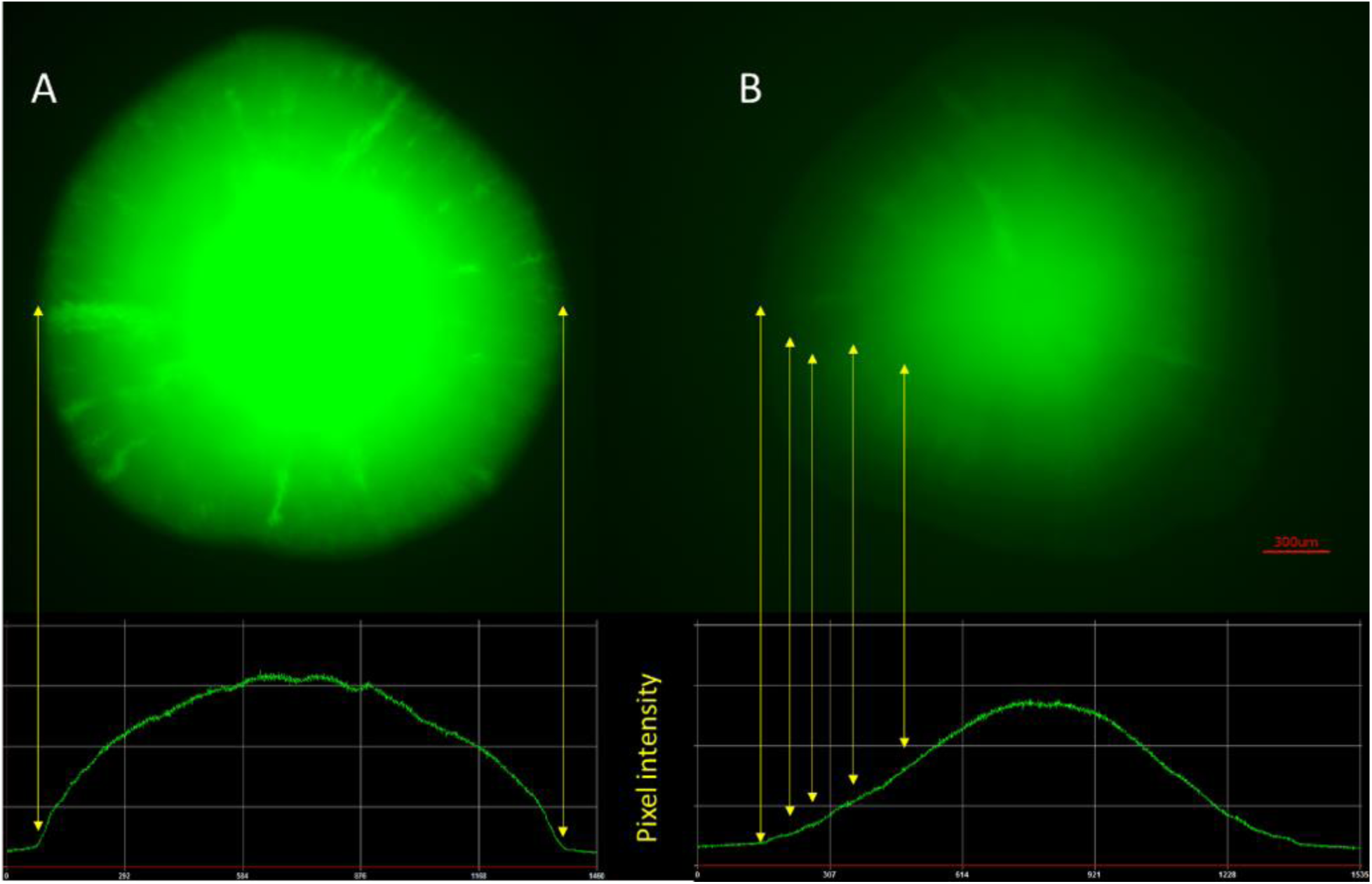
Microscopic picture of the 72h old *E. coli* C (A) and K12 (B) colonies containing pJEKd1571 grown at 37°C on LB Miller agar plates. Pixel intensity plots for each colony are shown below. Yellow arrows showed the colony borders and distinct *pcsrA* expression intensities.

### Location of IS3-like insertions in *E. coli* C genome and role of ISs in biofilm gene expression

Based on the *E. coli* C *pcsrA* promoter structure in comparison to the *pcsrA*-K12 (51) and its transcriptional activities, we concluded that the small 80-bp region containing the p4 and p5 promoters could not be solely responsible for the *csrA* transcription. Insertion sequences play a huge role in bacterial genome evolution (64). They can also insert upstream of a gene and activate its expression (65). Out of 177 genes that were unique to *E. coli C*, 55 encoded transposases (**S1 Tab.**).

Using BLAST, we found that the IS3-like sequence present in front of the *csrA* gene was present in 19 other locations throughout the genome (data not shown). Analyzing these locations, we found that in 12 cases the IS3 might drive the expression of downstream located genes (**Tab. 3**). One of the most striking observation was that the IS3-like sequence was located in front of an alternative sigma^70^ factor, which was not present in the K12 strain (**Tab. 3**). Based on the *pcsrA* expression, we concluded that a promoter located inside the IS3 drives permanent expression of the following genes. If that was the case, all those genes should have the same expression profile in different conditions, like in biofilm and planktonic cells. Based on our RNA-seq results, we found that in fact 8 out of 12 genes (regions) showed similar expression levels in both conditions (**Tab. 3**). However, the general expression level of gene B6N50_12000 encoding the alternative sigma^70^ factor was high, it was also highly overexpressed in biofilm (Diff.Expr.Log2 Ratio = 2.4; p value = 0.016) (**Tab. 3**). The presence of the constitutively expressed alternative sigma^70^ factor in the *E. coli* C can drive expression of the sigma^70^ promoters in a growth phase independent manner. As the remaining *E. coli* C *csrA* promoters P4 and P5 are sigma^70^ dependent promoters (51), it might explain their strong activity along all the cell growth phases. Further studies will be conducted to prove that hypothesis.

## Conclusions

As the bacterial genome undergoes a constant evolution and adaptation (66) and bacterial mobile elements are the most common mechanism of those processes (67, 68), one may ask why in this particular strain, unlike the other laboratory strains, the selection toward planktonic cells did not took place. There is no simple answer; however, we can speculate that as this strain was used for a proliferation of bacteriophages the fact that phages kill planktonic cells might reduce the selection toward free floating cells. The second hypothesis is that for bacteriophage research using the *E. coli* C, the ATCC recommends low-salt (0.5% NaCl) or no salt Nutrient (#139) broth medium. As we showed, the low-salt medium reduced bacterial stress and most likely reduced the level of genome rearrangements, keeping the natural properties for biofilm formation characteristic for the wild-type strains in this laboratory *E. coli* C strain.

Biofilms are the most prevalent form of bacterial life (9, 28) and as such have drawn significant attention from the scientific community over the past quarter century. However, only in 2018 did the number of biofilm related articles reach 24,000, based on a Google Scholar search. As in all other fields, biofilm research needs to develop and follow standard protocols and methods that can be used in different laboratories and give comparable results. Unfortunately, a standardized methodological approach to biofilm models has not been adopted, leading to a large disparity among testing conditions. This has made it almost impossible to compare data across multiple laboratories, leaving large gaps in the evidence (69). In our work, we described and characterized biofilm formation in the classic laboratory strain, *E. coli* C (2, 70). We have used that strain in our biofilm-related research for almost a decade and we would like to share it with the biofilm community and propose to use it as a model organism in *E. coli*-based biofilm-related research.

## Materials and Methods

### Bacterial strains and growth conditions

Bacterial strains are listed in Table 4. Strains were grown in M9 with glycerol medium or LB Miller, LB Lennox, or modified Lennox with 0.75% NaCl broth with appropriate antibiotics, kanamycin (50 µg/ml), gentamycin (10 µg/ml), and chloramphenicol (30 µg/ml).

**Table 1.** Sixty-nine unique genes in *E. coli* C genome

| Lp | Gene Name or Prokka group | Function - gene description |
| --- | --- | --- |
| 1 | mhpB_2 | 2,3-Dihydroxyphenylpropionate/2,3-dihydroxycinnamic acid 1,2-dioxygenase |
| 2 | yniC_1 | 2-Deoxyglucose-6-phosphate phosphatase |
| 3 | mhpC_2 | 2-Hydroxy-6-oxononadienedioate/2-hydroxy-6-oxononatrienedioate hydrolase |
| 4 | mhpD_2 | 2-Keto-4-pentenoate hydratase |
| 5 | mhpA_2 | 3-(3-Hydroxy-phenyl)propionate/3-hydroxycinnamic acid hydroxylase |
| 6 | mhpE_2 | 4-Hydroxy-2-oxovalerate aldolase |
| 7 | pcaK_2 | 4-Hydroxybenzoate transporter PcaK |
| 8 | mhpF_2 | Acetaldehyde dehydrogenase |
| 9 | group_2290 | Acetyltransferase (GNAT) family protein |
| 10 | group_2332 | Alpha/beta hydrolase family protein |
| 11 | csbX_1 | Alpha-ketoglutarate permease |
| 12 | csbX_2 | Alpha-ketoglutarate permease |
| 13 | group_2293 | Ankyrin repeats (3 copies) |
| 14 | group_2295 | Ankyrin repeats (3 copies) |
| 15 | group_2370 | Ankyrin repeats (3 copies) |
| 16 | clpC | ATP-dependent Clp protease ATP-binding subunit ClpC |
| 17 | ftsH3 | ATP-dependent zinc metalloprotease FtsH 3 |
| 18 | ftsH4 | ATP-dependent zinc metalloprotease FtsH 4 |
| 19 | group_2325 | Cell envelope integrity inner membrane protein TolA |
| 20 | bcsA_2 | Cellulose synthase catalytic subunit [UDP-forming] |
| 21 | wzzB_1 | Chain length determinant protein |
| 22 | group_2304 | Colanic acid exporter |
| 23 | dtpD_2 | Dipeptide permease D |
| 24 | group_337 | DNA-binding transcriptional regulator AraC |
| 25 | group_258 | Esterase YqiA |
| 26 | group_343 | Fructosamine kinase |
| 27 | group_2307 | GalNAc(5)-diNAcBac-PP-undecaprenol beta-1,3-glucosyltransferase |
| 28 | group_2365 | Glutathione-regulated potassium-efflux system protein KefC |
| 29 | ltrA_1 | Group II intron-encoded protein LtrA |
| 30 | ltrA_2 | Group II intron-encoded protein LtrA |
| 31 | ltrA_3 | Group II intron-encoded protein LtrA |
| 32 | ltrA_4 | Group II intron-encoded protein LtrA |
| 33 | ltrA_6 | Group II intron-encoded protein LtrA |
| 34 | dmlR_8 | HTH-type transcriptional regulator DmlR |
| 35 | group_2333 | HTH-type transcriptional regulator DmlR |
| 36 | hyfB_4 | Hydrogenase-4 component B |
| 37 | lacR_2 | Lactose phosphotransferase system repressor |
| 38 | tdh_2 | L-Threonine 3-dehydrogenase |
| 39 | malI_1 | Maltose regulon regulatory protein MalI |
| 40 | mtlK_2 | Mannitol 2-dehydrogenase |
| 41 | pglA | N,N'-Diacetylbaicillosaminy-diphospho-undecaprenol alpha-1,3-N-acetyl-galactosaminyltransferase |
| 42 | group_190 | Outer membrane usher protein HtrE precursor |
| 43 | group_240 | Periplasmic dipeptide transport protein precursor |
| 44 | group_2366 | Phosphate-starvation-inducible E |
| 45 | pduV_2 | Propanediol utilization protein PduV |
| 46 | nepI_2 | Purine ribonucleoside efflux pump NepI |
| 47 | group_2 | Putative deoxyribonuclease RhsC |
| 48 | group_49 | Putative fimbrial-like adhesin protein |
| 49 | argK_2 | Putative GTPase ArgK |
| 50 | group_330 | Putative HTH-type transcriptional regulator YbbH |
| 51 | group_168 | Putative lysophospholipase |
| 52 | group_2341 | Putative oxidoreductase |
| 53 | hhoB | Putative serine protease HhoB precursor |
| 54 | group_2281 | Recombination protein F |
| 55 | rbtD | Ribitol 2-dehydrogenase |
| 56 | group_2319 | Ribokinase |
| 57 | araB_1 | Ribulokinase |
| 58 | hspA_1 | Spore protein SP21 |
| 59 | hspA_2 | Spore protein SP21 |
| 60 | trxC_2 | Thioredoxin-2 |
| 61 | lsrR_1 | Transcriptional regulator LsrR |
| 62 | group_2327 | Type I restriction enzyme EcoKI subunit R |
| 63 | hsdR_1 | Type I restriction enzyme EcoR124II R protein |
| 64 | hsdR_2 | Type-I restriction enzyme R protein |
| 65 | group_2288 | Tyrosine recombinase XerD |
| 66 | group_2310 | UDP-glucose 4-epimerase |
| 67 | xylB_2 | Xylulose kinase |
| 68 | group_2360 | YfdX protein |
| 69 | group_2361 | YfdX protein |

**Table 2.** Top 40 of *E. coli* C differentially expressed genes in biofilm and planktonic mode of growth ranked by p value.

| Name | Function | Diff.Expr.Log2 Ratio | Dif.Expr. p value | Name | Function | Diff.Expr.Log2 Ratio | Dif.Expr. p value |
| --- | --- | --- | --- | --- | --- | --- | --- |
| B6N50_09285 | hypothetical protein CDS | 4.850395771 | 1.69E-39 | B6N50_13170 | trp operon leader peptide CDS | -5.43943394 | 4.17E-31 |
| B6N50_14580 | cold-shock protein CspG CDS | 3.557298531 | 4.41E-30 | B6N50_06090 | hypothetical protein CDS | -4.451200178 | 1.23E-16 |
| B6N50_11655 | cold-shock protein CspB CDS | 2.669746081 | 4.11E-17 | B6N50_03345 | threonine/serine transporter TdcC CDS | -3.469115448 | 3.92E-14 |
| B6N50_18720 | hypothetical protein CDS | 3.663331901 | 3.45E-15 | B6N50_06085 | thiol disulfide reductase thioredoxin CDS | -3.155139261 | 9.04E-14 |
| B6N50_18870 | hypothetical protein CDS | 5.888797737 | 7.60E-15 | B6N50_13175 | anthranilate synthase component I CDS | -4.59448805 | 1.48E-10 |
| B6N50_14575 | cold-shock protein CDS | 3.190816414 | 3.03E-14 | B6N50_03110 | tryptophan-specific transporter CDS | -3.207303792 | 3.06E-10 |
| B6N50_07425 | colicin V production protein CDS | 3.972852208 | 1.77E-13 | B6N50_18780 | fadE CDS | -2.605414693 | 3.45E-09 |
| B6N50_19625 | hypothetical protein CDS | 4.396004437 | 5.21E-13 | B6N50_03360 | reactive intermediate/imine deaminase CDS | -2.426031411 | 1.80E-08 |
| B6N50_04460 | 5-formyltetrahydrofolate cyclo-ligase CDS | 2.838038222 | 1.97E-12 | B6N50_15640 | inner membrane transport permease YbhR CDS | -2.420058315 | 2.65E-08 |
| B6N50_11790 | hypothetical protein CDS | 2.233953837 | 1.18E-11 | B6N50_01135 | copper-binding protein CDS | -3.646639286 | 2.81E-08 |
| B6N50_19130 | hypothetical protein CDS | 2.832250116 | 1.32E-11 | B6N50_17040 | cyclic amidohydrolase CDS | -4.009443561 | 3.10E-08 |
| B6N50_16480 | octanoyltransferase CDS | 2.970256489 | 1.97E-11 | B6N50_12480 | hypothetical protein CDS | -2.319595956 | 4.66E-08 |
| B6N50_18715 | hypothetical protein CDS | 3.010093216 | 5.09E-10 | B6N50_01230 | DUF305 domain-containing protein CDS | -2.471533306 | 9.48E-08 |
| B6N50_11650 | cold-shock protein CspF CDS | 3.047189907 | 1.58E-09 | B6N50_20670 | PTS trehalose transporter subunit IIBC CDS | -2.460088041 | 2.34E-07 |
| B6N50_21800 | hypothetical protein CDS | 3.244145509 | 2.86E-09 | B6N50_15645 | hypothetical protein CDS | -2.326096198 | 3.60E-07 |
| B6N50_10140 | hypothetical protein CDS | 2.407561788 | 7.37E-09 | B6N50_07515 | hypothetical protein CDS | -2.066303195 | 5.93E-07 |
| B6N50_16180 | putrescine-ornithine antiporter CDS | 2.814378464 | 7.54E-09 | B6N50_03340 | serine/threonine dehydratase CDS | -2.589176789 | 9.26E-07 |
| B6N50_02560 | acrEF/envCD operon transcriptional regulator CDS | 2.99746876 | 7.78E-09 | B6N50_22020 | hypothetical protein CDS | -1.892112356 | 1.11E-06 |
| B6N50_06220 | ferredoxin CDS | 3.430136055 | 9.64E-09 | B6N50_11925 | LsrR family transcriptional regulator CDS | -2.439904012 | 1.68E-06 |
| B6N50_14570 | cold-shock protein CDS | 2.881590809 | 3.60E-08 | B6N50_21220 | anaerobic C4-dicarboxylate transporter DcuA CDS | -1.646350417 | 1.78E-06 |
| B6N50_22715 | transporter CDS | 2.82399159 | 4.65E-08 | B6N50_12095 | glutamate decarboxylase CDS | -2.499993182 | 3.17E-06 |
| B6N50_19920 | 30S ribosomal protein S20 CDS | 1.941085309 | 6.11E-08 | B6N50_18960 | D-glycero-beta-D-manno-heptose 1,7-bisphosphate 7-phosphatase CDS | -2.011238834 | 3.58E-06 |
| B6N50_11725 | hypothetical protein CDS | 2.042056006 | 1.08E-07 | B6N50_20675 | glucohydrolase CDS | -2.390652037 | 3.90E-06 |
| B6N50_10135 | PhoP regulon feedback inhibition membrane protein MgrB | 2.818073792 | 1.41E-07 | B6N50_02360 | 50S ribosomal protein L30 CDS | -1.994964005 | 4.51E-06 |
| B6N50_20545 | hypothetical protein CDS | 2.123626973 | 2.36E-07 | B6N50_09330 | protein MtfA CDS | -2.19664491 | 4.91E-06 |
| B6N50_23295 | asparagine synthetase A CDS | 1.799864168 | 4.93E-07 | B6N50_21140 | fumarate reductase subunit C CDS | -2.811473378 | 5.46E-06 |
| B6N50_20610 | DNA-binding protein CDS | 2.193750313 | 4.99E-07 | B6N50_03355 | PFL-like enzyme TdcE CDS | -1.941660612 | 6.14E-06 |
| B6N50_14300 | transcriptional regulator CsgD CDS | 2.938430574 | 1.47E-06 | B6N50_01375 | acid stress chaperone HdeA CDS | -2.394334187 | 9.39E-06 |
| B6N50_04470 | cell division protein ZapA CDS | 1.635391947 | 1.64E-06 | B6N50_04060 | 8-amino-7-oxononanoate synthase CDS | -3.586412225 | 9.66E-06 |
| B6N50_05850 | hypothetical protein CDS | 2.359240446 | 1.64E-06 | B6N50_03350 | propionate kinase CDS | -2.252249192 | 1.15E-05 |
| B6N50_11345 | DNA-binding response regulator CDS | 1.889775289 | 2.01E-06 | B6N50_02275 | 50S ribosomal protein L4 CDS | -2.149717962 | 1.50E-05 |
| B6N50_11785 | sensor domain-containing diguanylate cyclase CDS | 1.86896654 | 2.43E-06 | B6N50_13195 | tryptophan synthase subunit alpha CDS | -2.656912572 | 1.77E-05 |
| B6N50_23245 | hypothetical protein CDS | 1.721769367 | 2.47E-06 | B6N50_06710 | hypothetical protein CDS | -1.739047636 | 1.95E-05 |
| B6N50_08970 | LPS O-antigen chain length determinant protein WzzB CDS | 1.839986113 | 3.18E-06 | B6N50_04225 | L-asparaginase 2 CDS | -1.58905668 | 3.32E-05 |
| B6N50_13505 | 4-diphosphocytidyl-2C-methyl-D-erythritol kinase CDS | 1.858003639 | 3.21E-06 | B6N50_22845 | fatty acid oxidation complex subunit alpha FadB CDS | -1.890631947 | 3.59E-05 |
| B6N50_08170 | bifunctional murein DD-endopeptidase/murein LD-carboxypeptidase | 1.72790582 | 4.36E-06 | B6N50_07440 | lysine/arginine/ornithine-binding periplasmic protein CDS | -2.2933939 | 3.64E-05 |
| B6N50_11695 | protein GnsB CDS | 1.610685956 | 5.02E-06 | B6N50_23470 | low affinity tryptophan permease CDS | -2.158586386 | 3.67E-05 |
| B6N50_16445 | ribosome silencing factor CDS | 1.691674447 | 6.54E-06 | B6N50_15500 | small protein MntS CDS | -1.558155635 | 3.80E-05 |
| B6N50_02975 | 50S ribosomal protein L21 CDS | 1.721186467 | 6.54E-06 | B6N50_03365 | L-serine dehydratase TdcG CDS | -1.680133634 | 3.90E-05 |
| B6N50_01245 | hypothetical protein CDS | 3.05358375 | 6.58E-06 | B6N50_17910 | transcriptional regulator CDS | -2.0922916 | 3.99E-05 |

**Table 3.** Genes located downstream of the IS3 like element and their differential expression in RNA-seq experiment.

|  | **Gene name** | **Function** | **Diff.Expr.Log2**<br>**Ratio**<br>**biofilm/planktonic** |
| --- | --- | --- | --- |
| 1. | B6N50_06210-15<br>(2 genes) | Unknown function and small toxic protein ShoB | 2.66; p = 0.018 |
| 2 | B6N50_07905-890 (4 genes) | Acetate CoA-transferase subunit alpha, acetate CoA-transferase subunit beta, short-chain fatty acids transporter and acetyl-CoA acetyltransferase | -2.55; p = 0.014 |
| 3 | B6N50_12005-12000 (2 genes) | Restriction endonuclease (pseudogene- missing start), RNA polymerase subunit sigma-70 (pseudogene- missing stop) | 2.4; p = 0.016 |
| 4 | B6N50_11960-55<br>(2 genes) | Hypothetical protein, hypothetical protein | 0.01; p = 0.9 |
| 5 | B6N50_05830 (1<br>gene) | Hypothetical protein | -0.49; p = 0.38 |
| 6 | B6N50_05870 (1<br>gene) | DEAD/DEAH box helicase | -0.05; p = 0.9 |
| 7 | B6N50_10145 (1<br>gene) | Hypothetical protein | 1.3; p = 0.19 |
| 8 | B6N50_11585 (1<br>gene) | Hypothetical protein | 0.48; p = 0.79 |
| 9 | B6N50_11780-85<br>(2 genes) | TIGR00156 family protein, sensor domain-containing diguanylate cyclase | 0.43; p = 0.54/2.07; p = 6.06E-05 |
| 10 | B6N50_11960 (1 gene) | Hypothetical protein | 0.01; p = 0.99 |
| 11 | B6N50_12235-260 (6 genes) | NarK family nitrate/nitrite MFS transporter, nitrate reductase subunit alpha, nitrate reductase subunit beta, nitrate reductase molybdenum cofactor assembly chaperone NarW, respiratory nitrate reductase subunit gamma, hypothetical protein | 0.11; p = 0.96 |
| 12 | B6N50_19300-320 (5 genes) | Outer membrane usher protein, fimbrial protein, fimbrial protein StaE, fimbrial protein StaF, hypothetical protein | 0.4; p = 0.54/ staE 1.5; p = 3.8 |

**Table 4.** Bacterial strains used in this work.

| Strain | Genotype | Source |
| --- | --- | --- |
| *E. coli* C | WT | Holly A. Wichman, James Bull |
| *E. coli* K12 MG1655 | WT | Lab collection |
| *E. coli* Crooks | WT | Lonnie Ingram |
| *E. coli* B | WT | ATCC |
| *E. coli* W | WT | ATCC |
| *E.coli* K12 *csrA* | *csrA*::mini-Tn5 Km<sup>R</sup> | Tony Romeo |
| *E. coli* K12 MG1655 EC100 | F<sup>-</sup>*mcrA* Δ(*mrr-hsdRMS-mcrBC*) φ80*dlacZ*ΔM15 Δ*lacX74 recA1 endA1 araD139* Δ(*ara, leu*)7697 *galU galK λ-rpsL nupG*. | Lucigen |
| **Plasmids** |  |  |
| pBBR1MCS-5 | Broad host range mobilizable plasmid, Gm<sup>R</sup> | (52) |
| pAG136 | pET28B –EGFP-YFAST, Km<sup>R</sup> promoter probe vector | (62) |
| pProbe GFP[ASV] | Short-life GFP promoter probe vector, Km<sup>R</sup> | (63) |
| pJEKd1750 | 1638-bp fragment with *E.coli* C IS3-*csrA* promoter in BglII/XbaI sites of pAG136, Km<sup>R</sup> | This work |
| pJEKd1751 | 1638-bp fragment with *E. coli* C IS3-*csrA* promoter in SmaI site of pProbe GFP[ASV], Km<sup>R</sup> | This work |
| pJEK718 | *csrA* gene driven by the *plac* promoter in pBBR1MCS-5, Gm<sup>R</sup> | This work |
| pJEK786 | *csrA* gene driven by the *pcat*<br>promoter in pBBR1MCS-5,<br>Gm<sup>R</sup> Cm<sup>R</sup> | This work |

**Abbreviations:** Km<sup>R</sup>, Gm<sup>R</sup>, Cm<sup>R</sup> – kanamycin, gentamycin, and chloramphenicol resistance.

## All methods are described in Supplementary Materials

### Data availability

The RNA-seq raw sequencing data have been deposited in the GEO repository with accession code GSE125110. All other relevant data are available in this article and its Supplementary Information files. The complete genome of *E. coli* C has been deposited in GenBank under accession no. CP020543.1.

## ACKNOWLEDGMENTS

This research was supported by the CGS and CAMP at Drexel University. Thanks to Drs. Eva M. Top, Holly A. Wichman (University of Idaho), James Bull (University of Texas) for the *E. coli* C strain. Drs. Lonnie Ingram, Tony Romeo (University of Florida), Steven E. Lindow (University of California), and Arnaud Gautier (PASTEUR UMR8640 ENS-PSL University / Sorbonne Université / CNRS) for their strains and plasmids. We thank Drs. Brian Wighdal and Michael Nonnemacher for using the ImageQuant TL software. We thank Jocelyn Hammond for proofreading and comments.

---

**S1 Fig. Neighbor joining tree based on gene presence/absence within five *E. coli* strains.**

**S2 Fig. Genome view of (A) K12 LPS (*waa*) regions blast results with *E. coli* C genome and (B) colonic acid (*wca*) region in *E. coli* C.** Red double-headed arrow shows deleted region in strain C. Green arrow indicates a long operon like stretch of 35 genes with IS3 insertions in *wzzB* and B6N50_08940 genes (black double-headed arrows).

**S3 Fig. Genome view of K12 *fim* region blast results with *E. coli* C genome.** Red double-headed arrow shows deleted region in strain C.

**S4 Fig. Genome view of K12 *yhc* region blast results with *E. coli* C genome (A) and the *E. coli* C *yad* region with two IS insertions (black arrows).** Deletion of IS5 in *E. coli* C *yhcE* gene is highlighted.

**S5 Fig. Genome view of K12 (upper) and C strain (lower) *csg* region blast results with *E. coli* C genome.** Red double-headed arrow shows region replaced by IS5 in strain C.

**S6 Fig. PCR amplification of the *csrA* gene from *E. coli* C and K12 strains.**

**S7 Fig. Biofilm formation by *E. coli* C and K12 *csrA* mutant strains on (A) microscope slides (LB medium-72h) and (B) 96-well plates (LB Miller broth 37°C; 24h).**

**S8 Fig. PCR amplification of the *alaS-csrA* intergenic region from *E. coli* C and K12 strains.**

**S8 Fig. Differences in relative *pcsrA* promoter activity between *E. coli* C and K12 strains grown in LB Miller broth at 24°C and 37°C (250 rpm).** Cell densities (OD_600_) and fluorescence (480 nm Ex./520 nm Em.) were measured over the time course to show the relative promoter activity in each strain and condition. The graph represents the ratios between these activities in *E. coli* C and K12 strains at the specific time points.

**S10 Fig. Cell aggregation of *E. coli* C and K12 carrying the pJEKd1750 plasmid in overnight culture grown at 37°C in LB Miller broth on shaker at 250 rpm.** Ratio of planktonic cells to total cells measured as OD_600_.

**S1 Fig.**
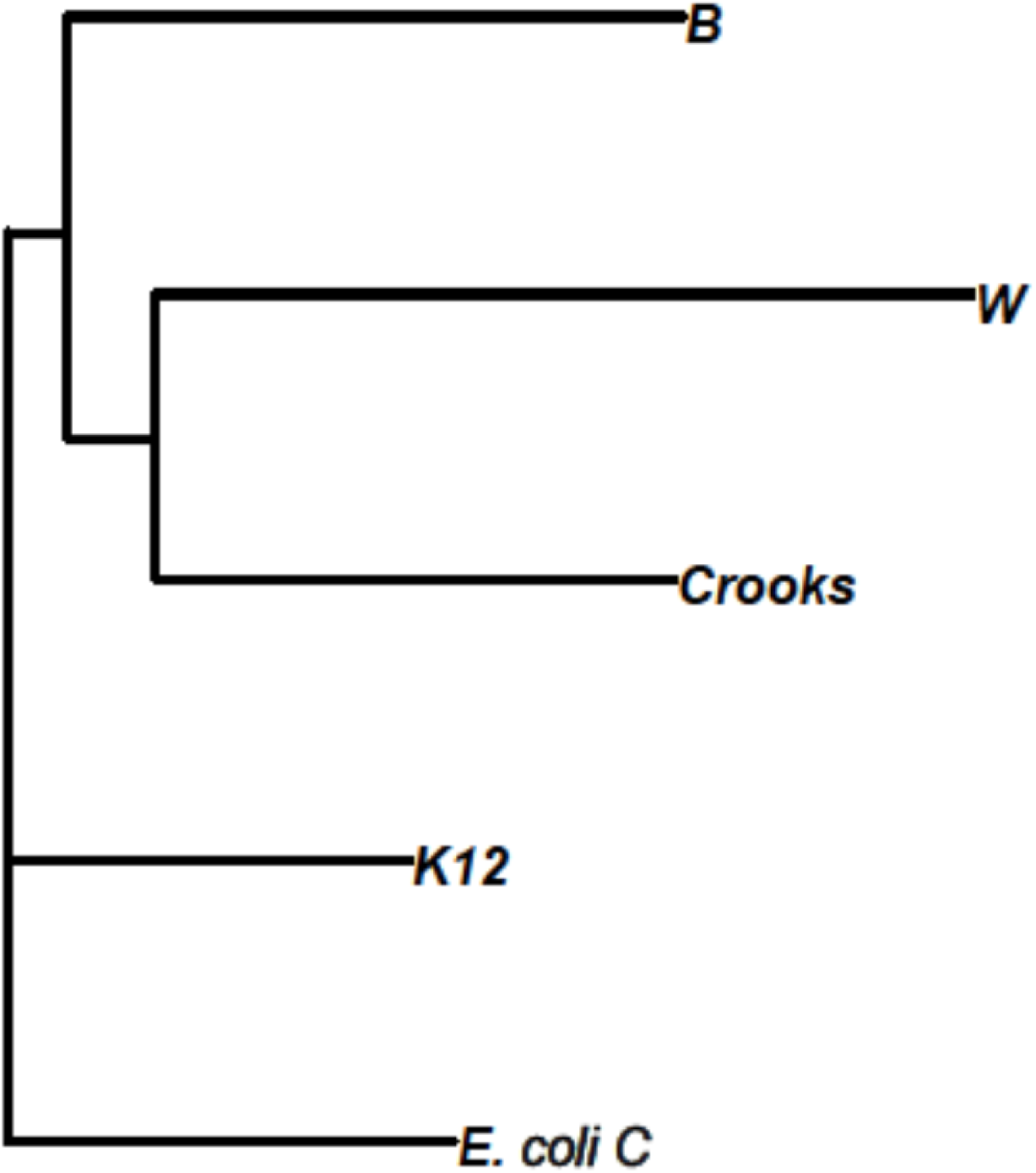
Neighbor joining tree based on gene presence/absence within five *E. coli* strains.

**S2 Fig.**
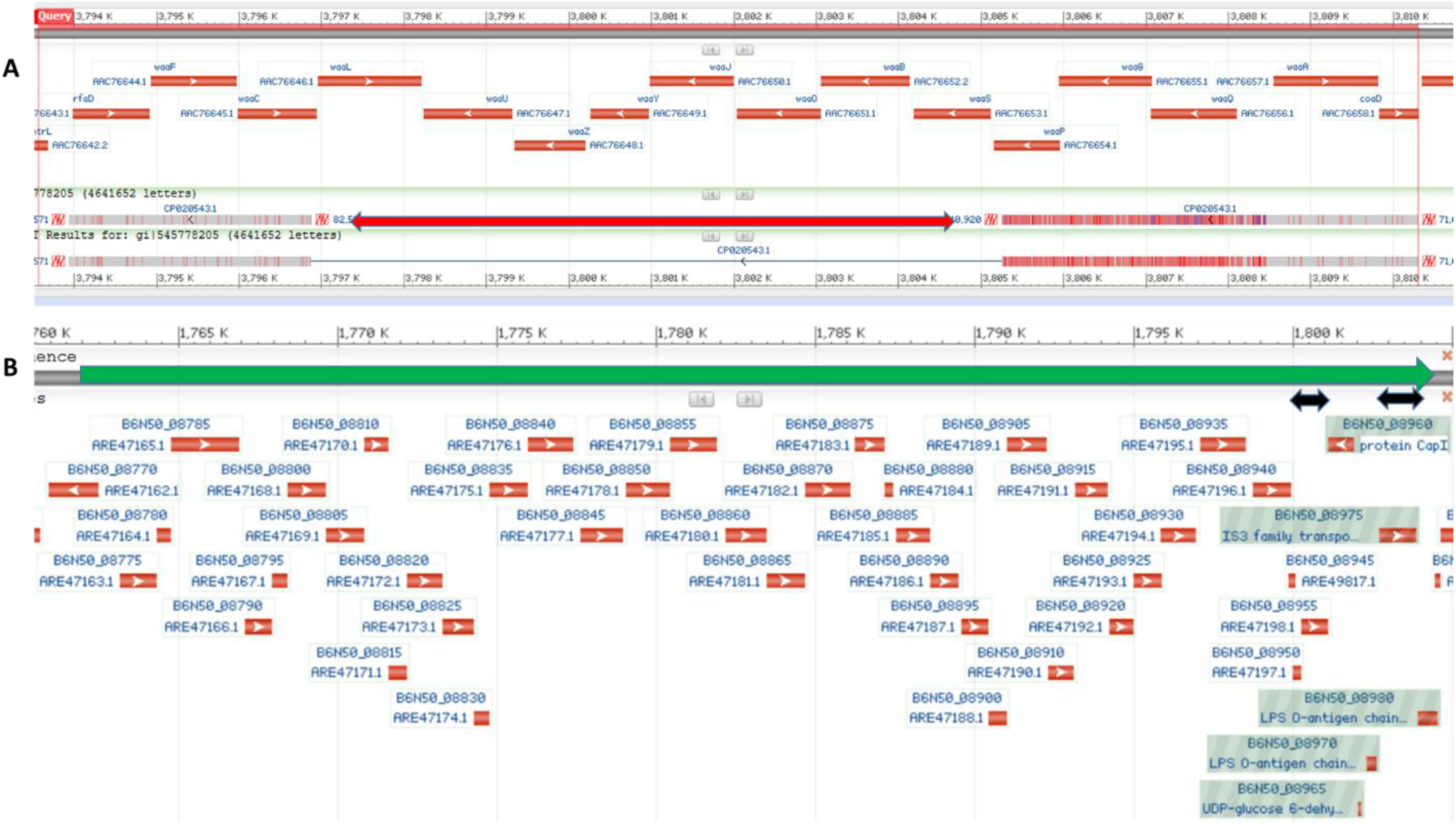
Genome view of (A) K12 LPS (*waa*) regions blast results with *E. coli* C genome and (B) colonic acid (*wca*) region in *E. coli* C. Red double-headed arrow shows deleted region in strain C. Green arrow indicates a long operon like stretch of 35 genes with IS3 insertions in *wzzB* and B6N50_08940 genes (black double-headed arrows).

**S3 Fig.**
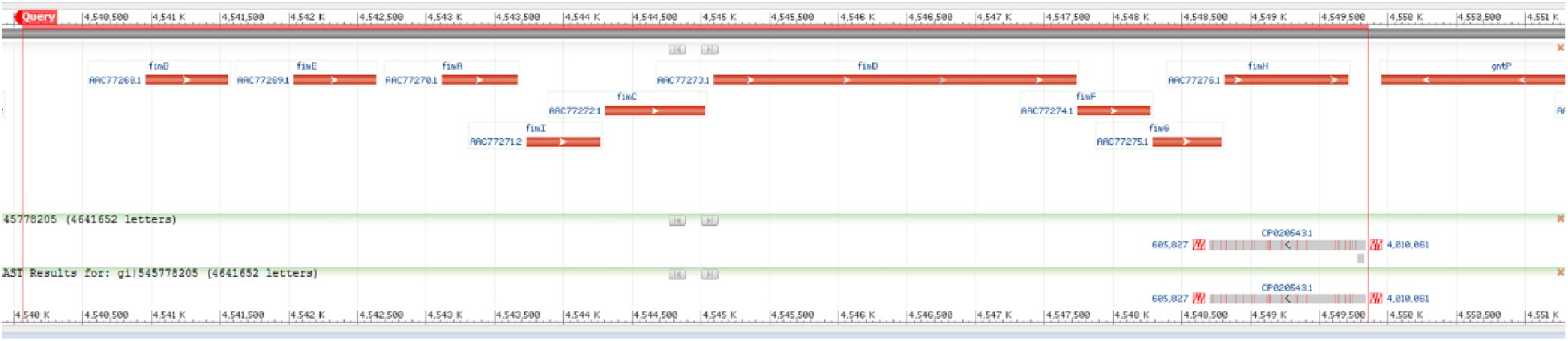
Genome view of K12 *fim* region blast results with *E. coli* C genome. Red double-headed arrow shows deleted region in strain C.

**S4 Fig.**
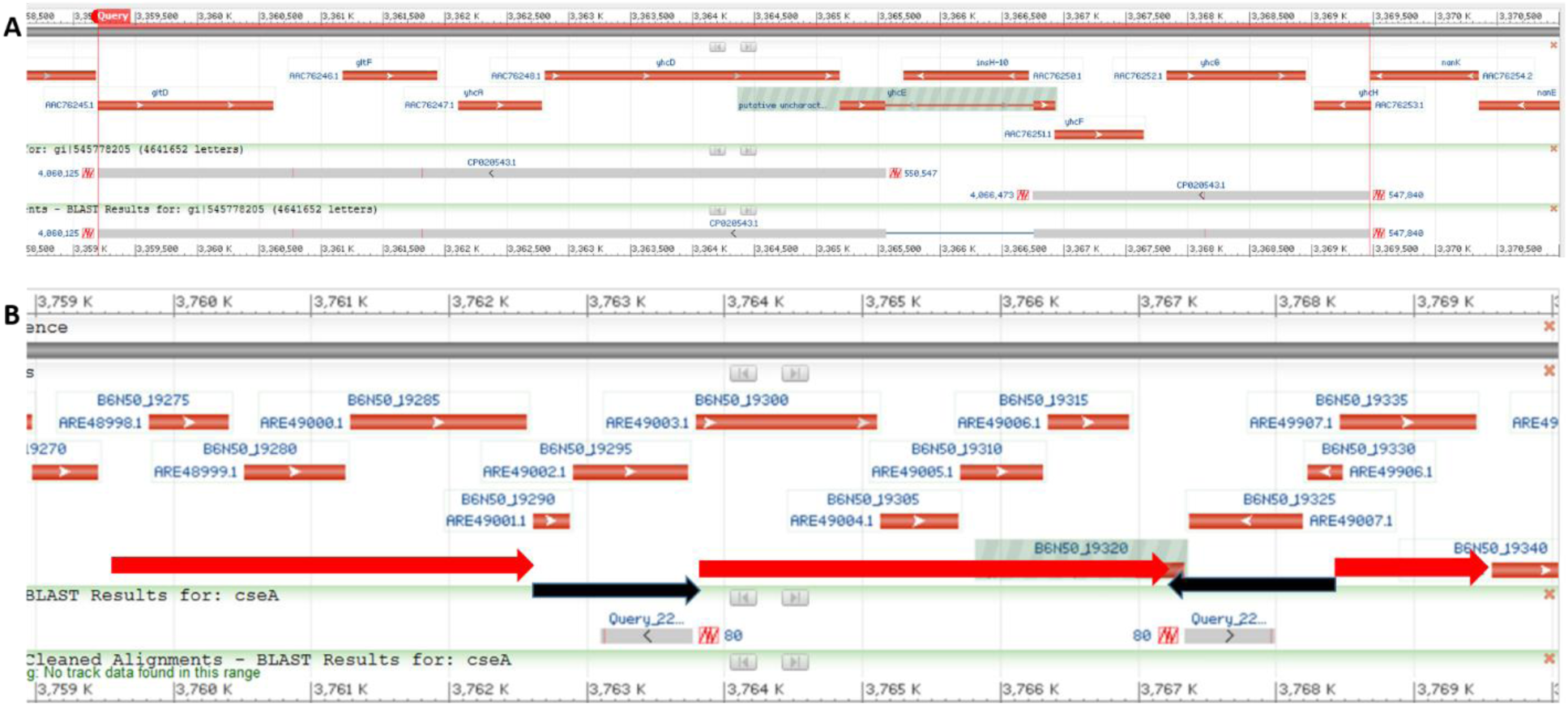
Genome view of K12 *yhc* region blast results with *E. coli* C genome (A) and the *E. coli* C *yad* region (B) with two IS insertions (black arrows). Deletion of IS5 in *E. coli* C *yhcE* gene is highlighted.

**S5 Fig.**
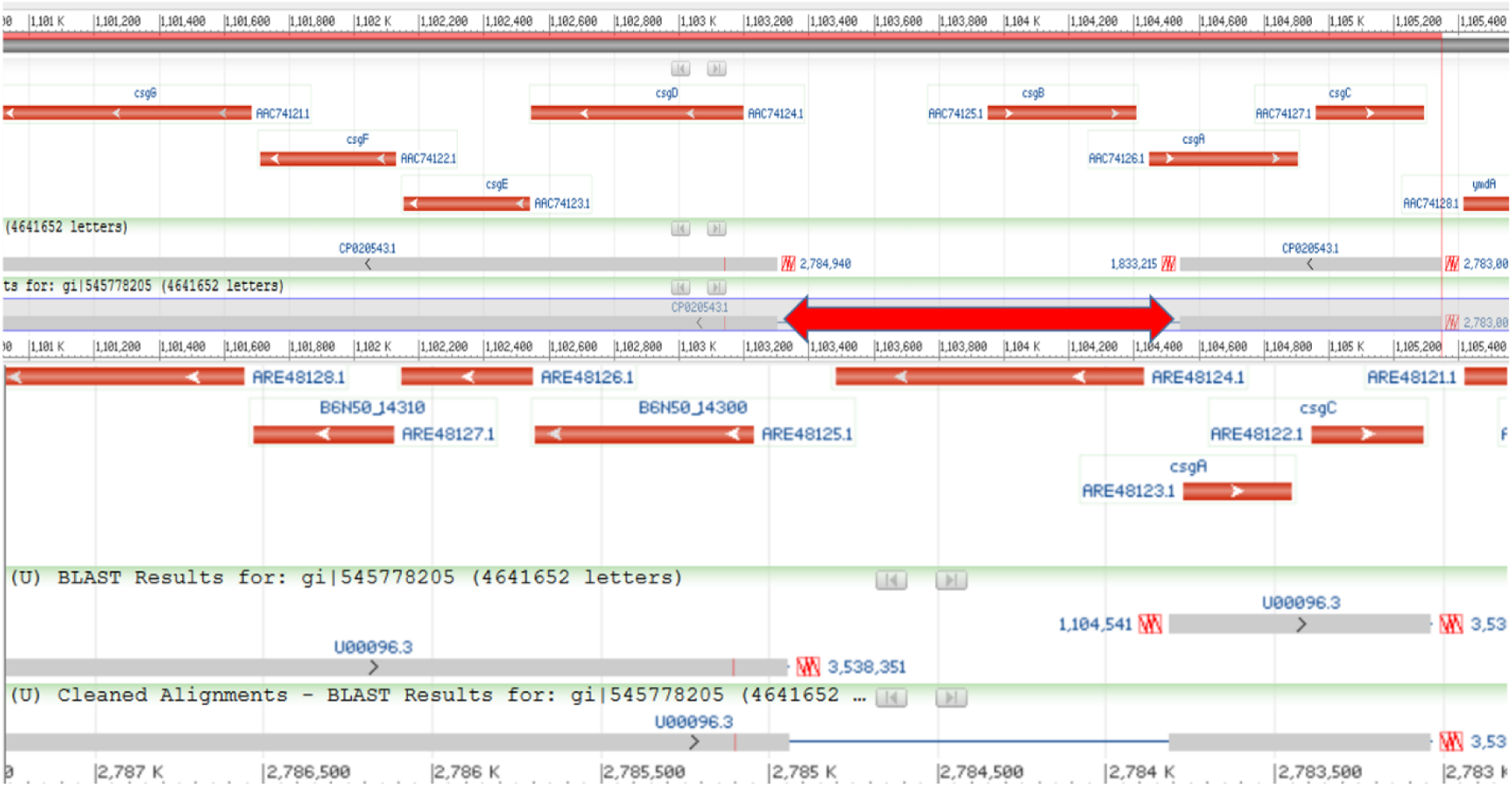
Genome view of K12 (upper) and C strain (lower) *csg* region blast results with *E. coli* C genome. Red double-headed arrow shows region replaced by IS5 in strain C.

**S6 Fig.**
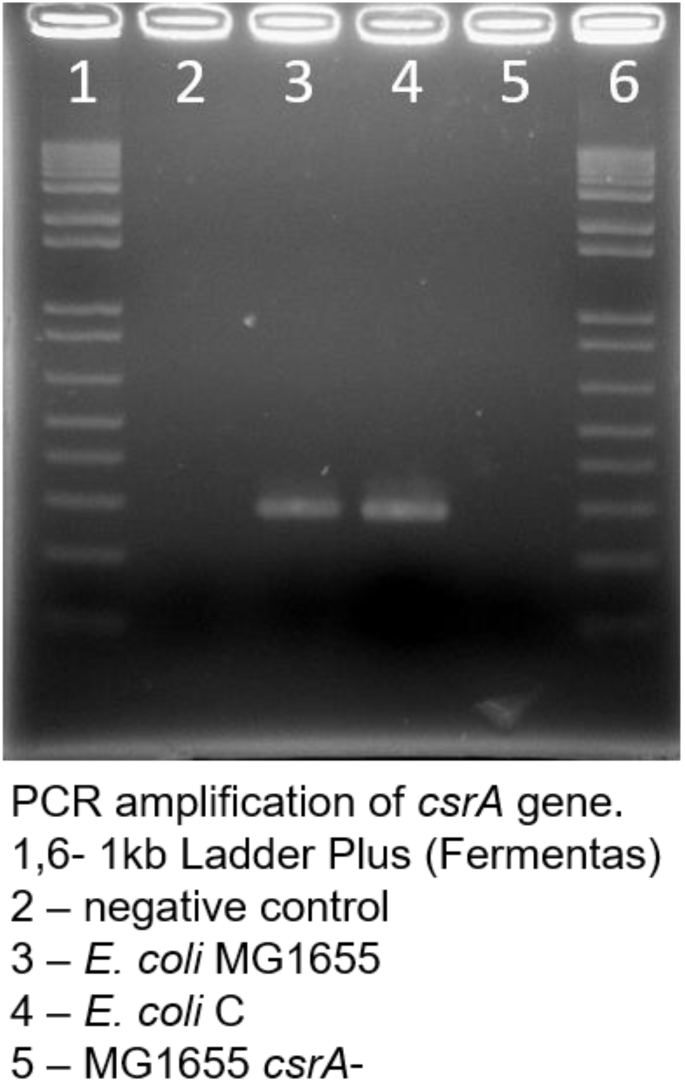
PCR amplification of the *csrA* gene from *E. coli* C and K12 strains.

**S7 Fig.**
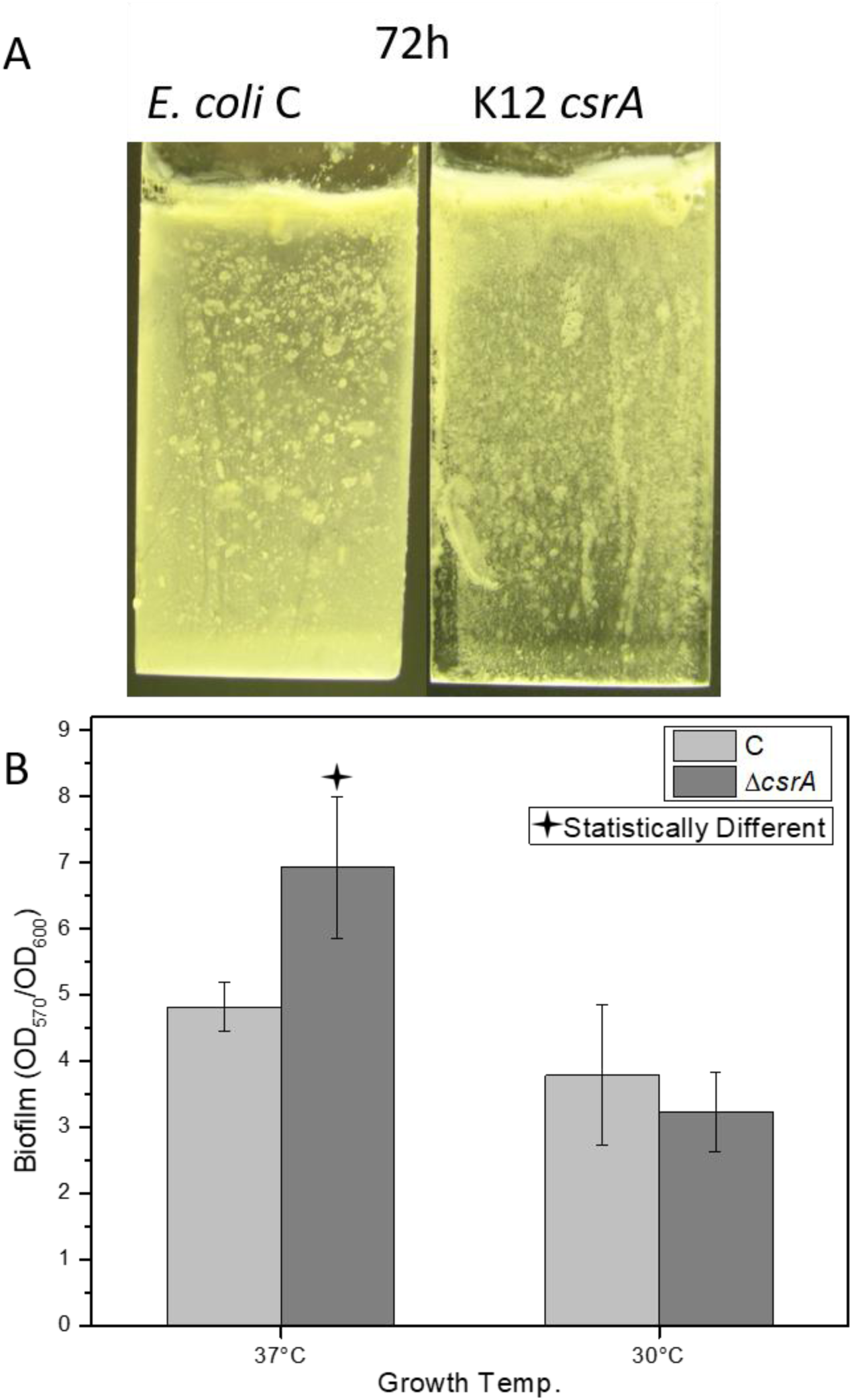
Biofilm formation by *E. coli* C and K12 *csrA* mutant strains on (A) microscope slides (LB medium-72h) and (B) 96-well plates (LB Miller broth 37°C; 24h).

**S8 Fig.**
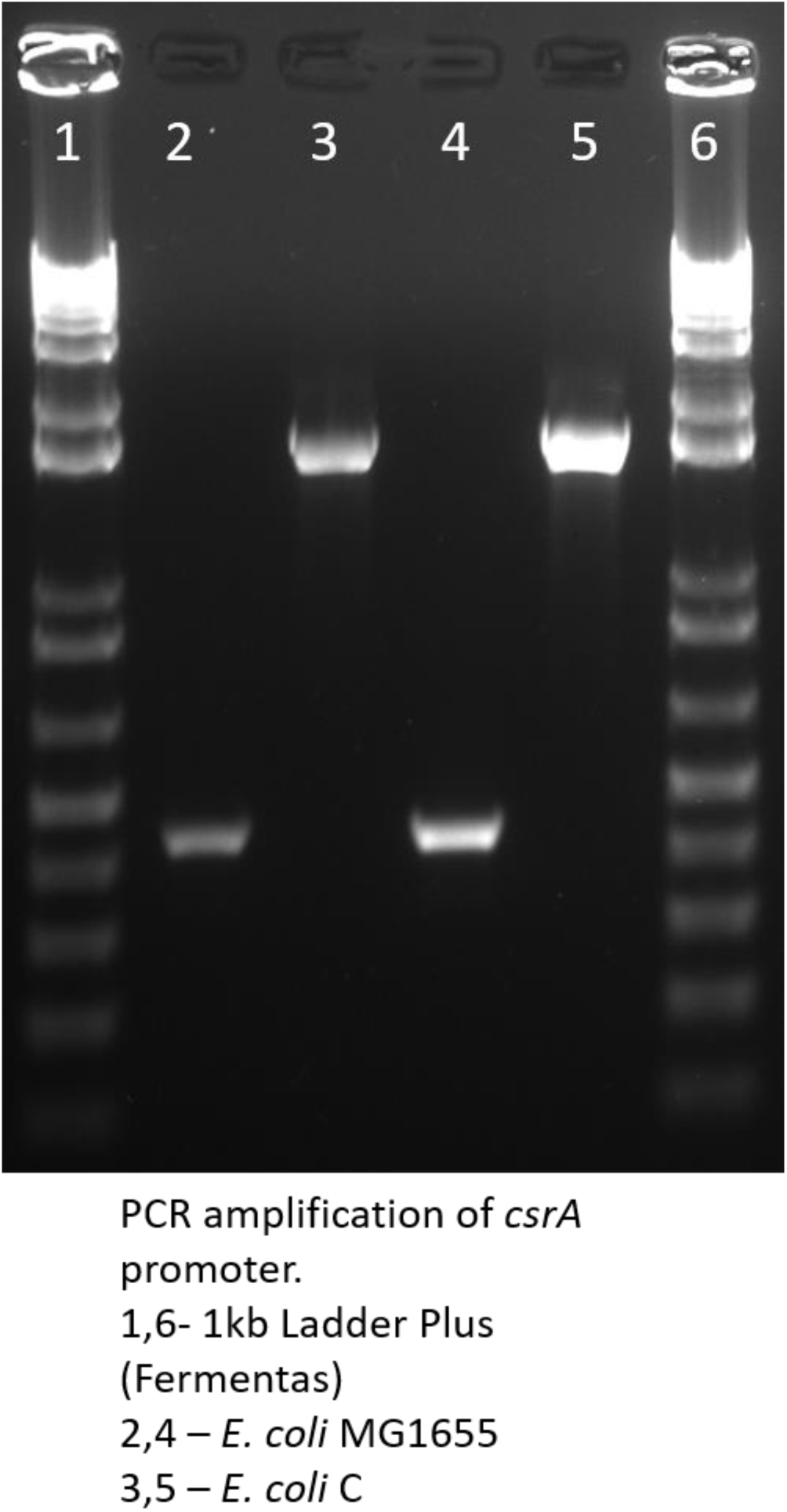
PCR amplification of the *alaS-csrA* intergenic region from *E. coli* C and K12 strains.

**S9 Fig.**
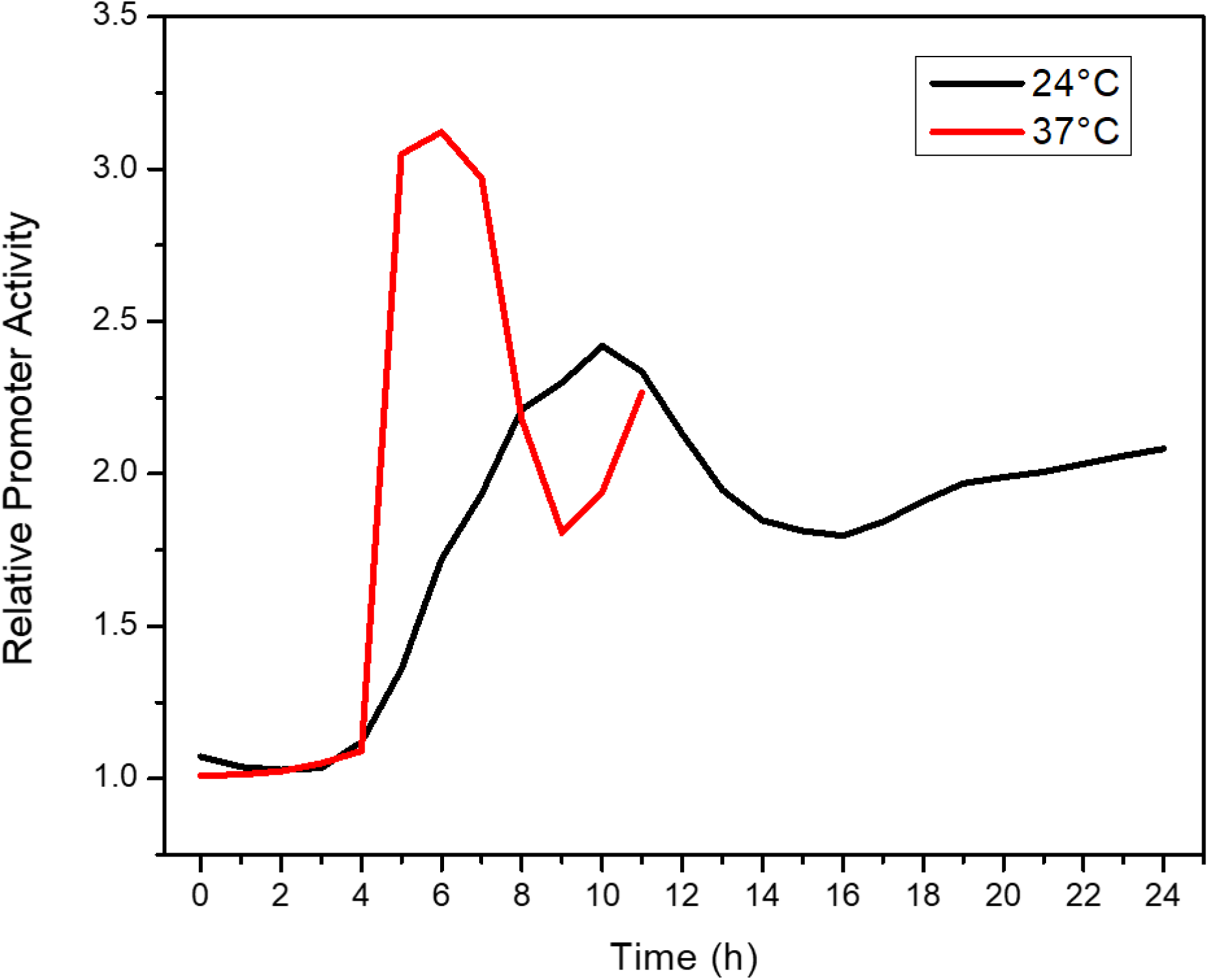
Differences in relative *pcsrA* promoter activity between *E. coli* C and K12 strains grown in LB Miller broth at 24°C and 37°C (250 rpm). Cell densities (OD_600_) and fluorescence (480 nm Ex./520 nm Em.) were measured over the time course to show the relative promoter activity in each strain and condition. The graph represents the ratios between these activities in *E. coli* C and K12 strains at the specific time points.

**S10 Fig.**
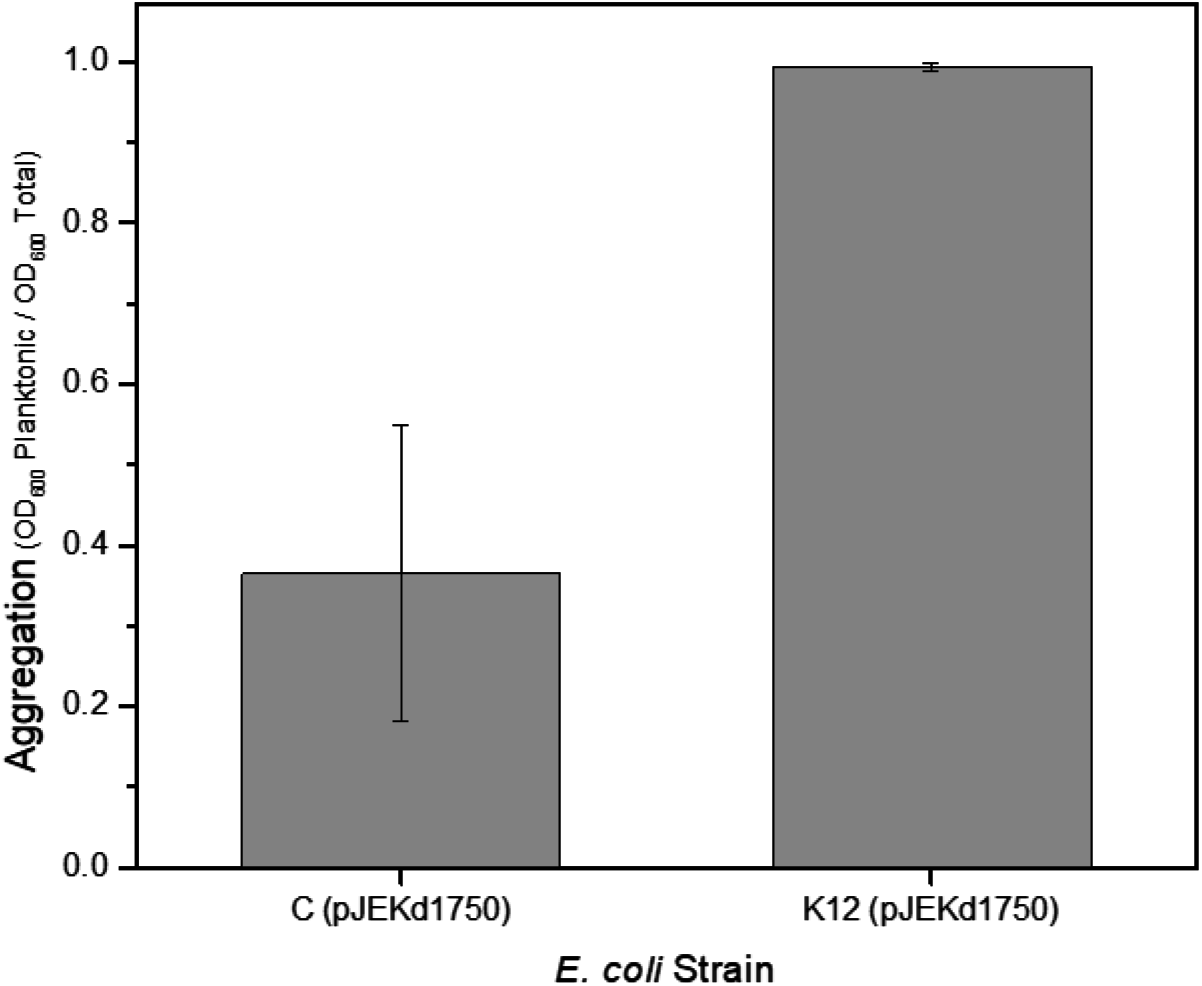
Cell aggregation of *E. coli* C and K12 carrying the pJEKd1750 plasmid in overnight culture grown at 37°C in LB Miller broth on shaker at 250 rpm. Ratio of planktonic cells to total cells measured as OD_600_.

## Supplementary Materials

### Methods

#### Biofilm assays

Biofilms on microscope slide were grown as described previously (1). For biofilm formation on a polystyrene surface, flat-bottom 96-well microtiter plates (Corning Inc.) were used (2). *E. coli* overnight cultures were diluted 1:40 in fresh medium, and 150-μL aliquots were dispensed into wells. After 24 hours of incubation (37°C), cell density was measured (OD_600_) using a plate reader, and 30 μL of Gram Crystal Violet (Remel) was applied for staining for 1 hour. Plates were washed with water and air dried, and crystal violet was solubilized with an ethanol-acetone (4:1) solution. The OD_570_ was determined from this solution, and the biofilm amount was calculated as the ratio of OD_570_ to OD_600_ (Krol et al. 2014).

#### Construction of CsrA overexpressing strain

A 277-bp DNA fragment containing the csrA gene was amplified using csrAF-aaa GAATTCGTAATACGACTCACTATAGGGTTTC csrAR – aaaGAATTCTTTGAGGGTGCGTCTCACCGATAAAG primers. This fragment was cloned directly into the *Eco*RI site of the pBBR1MCS-5 vector (3). Sequence orientation was verified by DNA sequencing and the correct clone with *csrA* gene downstream of the *plac* promoter was named pJEK718. To express the *csrA* gene with a constitutive *pcat* (chloramphenicol) promoter, a PCR amplified *cat* gene (870 bp, catF-aaaGATCCTGGTGTCCCTGTTGATACCGGGAA; cat-R-aaa GGATCCCCCAGGCGTTTAAGGGCACCAATAAC) was cloned in the *Bam*HI site of one of the clones which carried the *csrA* gene in the orientation opposite to the *plac* promoter in the pBBR1MCS-5 vector. Selection for Cm-resistant clones ensured the promoter activity and the correct orientation was verified by PCR with catF/csrAR primers and DNA sequencing. The correct plasmid was named pJEK786. Plasmids were introduced into the *E. coli* C strain by TSS transformation (4).

#### Confirmation of IS3 insertion and construction of GFP reporter fusions

PCR fragments containing *csrA* promoter were amplified using pcsrA aaaagatctCTGATTGCAGGCGTATCTAAGG and pcsrAR aaatctagaAAAGATTAAAAGAGTCGGGTCTCTCTGTATCC primer pair from both *E. coli* K12 and C strains and cloned into the BglII/XbaI site of the pAG136 plasmid (5).or the *Sma*I site of the pPROBE-GFP[LVA] promoter probe vector (6). All constructs were verified by DNA sequencing. Plasmids were introduced into both the *E. coli* K12 and C strains by a TSS transformation (4). GFP activity (OD_480-520_) was measured using Bio-Tek Synergy HT (BioTek) or Tecan InfiniteM200 Pro (Tecan) plate readers and normalized to the optical density of the culture (OD_600_), yielding relative fluorescence units (RFU; OD_480-520_/OD_600_). For quantification of promoter activities in late stationary phase, single colonies were inoculated into 5 mL of LB broth, vortexed and 5 µL of cell suspension was spotted on LB Miller agar plates. Plates were incubated at 30°C or 37°C. At the specific time points, plates were scanned with a Typhoon 9400 Variable Mode Imager using 532/526-nm excitation/emission wavelengths (GE Healthcare). Scans were analyzed using the ImageQuant TL software (GE Healthcare). Student T-test was used to compare results and check statistical significance. Fluorescence microscopy was done with a Keyence BZ-X710 All-in One Fluorescence microscope (Keyence).

#### Cell aggregation and precipitation experiments

*E. coli* strains were grown in LB Miller broth at 30°C in shaking conditions (250 rpm). One milliliter of the culture was transferred to standard polypropylene spectrophotometer cuvettes to measure planktonic cells densities (OD_600_). Remaining cultures were vortexed ∼1 minute and 1 mL was aliquoted into cuvettes to measure the total cell densities (OD_600_). Aggregation was calculated as a ratio of planktonic to total cell density. For the precipitation experiment, overnight cultures were vortexed ∼1 minute and 1 mL was aliquoted into standard polypropylene spectrophotometer cuvettes and capped. Cuvettes were incubated statically at 12°C, 24°C (room temperature), and 37°C. Cell densities were measured every hour by measuring OD_600_.

#### DNA sequencing and sequence analyses

DNA for sequencing was isolated using the Qiagen Blood and Tissue DNA Isolation Kit. Genomic DNA was mechanically sheared using a Covaris g-TUBE. The SMRTbell template preparation kit 1.0 (Pacific Biosciences, Menlo Park, CA, USA) was used according to the PacBio standard protocol (10-kb template preparation using the BluePippin size-selection system Sage Science). After SMRTbell preparation and polymerase binding, the libraries were loaded on SMRTcells via magbead loading and run on a PacBio RS II instrument (Pacific Biosciences) using a C4 chemistry. DNA sequence data were assembled by the HGAP Assembly 2 and annotated by Prokka or NCBI’s Prokaryotic Genome Automatic Annotation Pipeline (PGAAP)(7). *E. coli* C, K12, B, W, and Crook genomes were analyzed by Roary, Mauve and Geneious R11.

#### Optical mapping - high molecular weight DNA extraction

Cells from overnight culture were washed with PBS, resuspended in cell resuspension buffer, and embedded into low-melting-point agarose gel plugs (BioRad #170-3592, Hercules, CA, USA). Plugs were incubated with lysis buffer and proteinase K for 4h at 50°C. Plugs were washed, melted, and solubilized with GELase (Epicentre, Madison, WI, USA). Purified DNA was subjected to 4h of drop-dialysis and DNA concentration was determined using Quant-iTdsDNA Assay Kit (Invitrogen/Molecular Probes, Carlsbad, CA, USA). DNA quality was assessed with pulsed-field gel electrophoresis. High molecular weight DNA was labeled according to commercial protocols with the IrysPrep Reagent Kit (Bionano Genomics). Roughly 300 ng of purified genomic DNA was nicked with 7 U of nicking endonuclease Nt.BspQI (NEB) at 37°C for 2h in NEB Buffer 3. Nicked DNA was labeled with a fluorescent-dUTP nucleotide analog using Taq polymerase (NEB) for 1h at 72°C. Nicks were repaired with Taq ligase (NEB) in the presence of dNTPs. The backbone of fluorescently labeled DNA was stained with YOYO-1 (Invitrogen). Labeled DNA molecules entered nanochannel arrays of an IrysChip (Bionano Genomics) via automated electrophoresis. Molecules were linearized in the nanochannel arrays and imaged. An in-house image detection software detected the stained DNA backbone and locations of fluorescent labels across each molecule. The set of label locations within each molecule defined the single-molecule maps. The *E. coli strain C* reference sequence was *in silico* nicked with Nt.BspQI. Raw single-molecule maps were filtered by minimum length of 150 kbp. Molecule maps were aligned to the *E. coli* reference map with OMBlast. OMBlast is an optical mapping alignment tool using a seed- and-extend approach and allows split-mapping (8). Alignments were performed with the OMBlastMapper module (version 1.4a) using the following parameters: --writeunmap false -- optresoutformat 2 --falselimit 8 --maxalignitem 2 --minconf 0. Molecule maps with partial alignments to regions flanking the putative insertion breakpoint coordinates were extracted from the alignment output file. Molecule maps were manually inspected for label matches in segments 5′ and 3′ to the putative inverted region and into the inversion. The non-aligned segments of these maps, which extended into the inverted region with label matches to the opposing side in a reverse fashion, were retained.

#### Biofilm gene expression profiling

Total RNA was extracted from 5-day-old slide biofilms, which were transferred to fresh LB Miller broth daily. As planktonic cells, we used cells detached from the biofilm from the last, 5^th^-day passage. TRIzol method (Invitrogen) extracted RNAs were processed with Illumina’s Ribozero bacteria kit to deplete rRNA; then Illumina’s TruSeq stranded RNA LT kit was used for library construction. The libraries were sequenced with the Illumina NextSeq 500 and a high output 75 cycle v2 flow cell. Reads were assembled to the *E. coli* C genome and analyzed by DEseq3 (9, 10) and Geneious R11 software. Volcano plot and its statistics were generated by the Geneious R11 software.

#### Statistical analysis

Statistical analysis was carried out in the R computing environment and in Graphpad. Relevant statistical information is included in the methods for each experiment. Error bars show standard deviation from the mean. Asterisk representing statistical significance p < 0.05.

